# A Fragment-based approach to assess the ligandability of ArgB, ArgC, ArgD and ArgF in the L-arginine biosynthetic pathway of *Mycobacterium tuberculosis*

**DOI:** 10.1101/2021.03.12.435067

**Authors:** Pooja Gupta, Sherine E. Thomas, James Cory-Wright, Víctor Sebastián-Pérez, Ailidh Burgess, Emma Cattermole, Clio Meghir, Chris Abell, Anthony G. Coyne, William R. Jacobs, Tom L. Blundell, Sangeeta Tiwari, Vítor Mendes

**Author notes:** Current affiliation: MRC-Laboratory of Molecular Biology, Molecular Immunity Unit, Francis Crick Ave, Cambridge, CB2 0QH, UK. To whom correspondence should be addressed, Vitor Mendes, Sangeeta Tiwari.

## Abstract

The L-arginine biosynthesis pathway consists of eight enzymes that catalyse the conversion of L-glutamate to L-arginine, appears to be attractive target for anti-Tuberculosis (TB) drug discovery. Starvation of *M.* tuberculosis deleted for either *argB* or *argF* genes led to rapid sterilization of these strains in mice while Chemical inhibition of ArgJ with Pranlukast was also found to clear chronic *M. tuberculosis* infection in animal models. In this work, the ligandability of four enzymes of the pathway ArgB, ArgC, ArgD and ArgF is explored using a fragment-based approach. We reveal several hits for these enzymes validated with biochemical and biophysical assays, and X-ray crystallographic data, which in the case of ArgB were further confirmed to have on-target activity against *M. tuberculosis*. These results demonstrate the potential of more enzymes in this pathway to be targeted with dedicated drug discovery programmes.

## 1. Introduction

Despite the availability of effective chemotherapy, tuberculosis (TB) remains a leading infectious cause of morbidity and mortality worldwide. In 2019, an estimated 1.2 million deaths were caused by TB, and an additional 208,000 were a result of HIV-TB co-infection (1). Simultaneously, the existing multidrug treatment regimen has a success rate of 85% in drug-sensitive TB cases (in the 2018 cohort), drug toxicity, a long treatment duration, and resulting patient non-compliance, as well as incompatibility with antiretroviral therapy all compromise its effectiveness. Alarmingly, the emergence of multi-drug resistant (MDR) and extensively-drug-resistant (XDR) strains of *Mycobacterium tuberculosis* has further undermined the efficacy of current antitubercular therapy: only 57% of MDR cases were successfully treated worldwide in the 2017 cohort. New antitubercular agents are therefore urgently required and novel chemical scaffolds and mechanisms of action must be identified that can shorten therapy and circumvent development of drug resistance. While many drugs can be bacteriocidal, *M. tuberculosis* has the ability to generate subpopulations that enter into a persister state making them phenotypically drug resistant (2). The consideration of preventing persister formation or killing persisters needs to be addressed in future drug discovery campaigns against *M.tuberculosis*.

*M. tuberculosis,* like the leprosy bacillus, has retained its ability to make all 20 amino acids and most vitamins. This retention of these biosynthetic genes reflect a evolutionary pressure suggesting the pathogenic mycobacteria have chosen not to obtain amino acids or many vitamins from the host and has thus been described as an autarkic lifestyle (3). However, not all amino acid auxotrophies behave the same. Several amino acid auxotrophs were found to have attenuated virulence inside host organisms, suggesting that while enzymes in amino acid biosynthetic pathways are essential *in vitro*, the pathogen can scavenge amino acids (albeit insufficiently) inside the host and survive (4–9). However, it was shown that methionine and arginine auxotrophs of *M. tuberculosis* are rapidly sterilised in both immunocompetent and immunodeficient (SCID) mice without the appearance of suppressor/bypass mutants (3, 10). Despite the presence of two arginine transporters in *M tuberculosis* (11, 12) and sufficiently high serum concentrations of arginine in the host (13), the virulence of Δ*argB* or Δ*argF* mutants is entirely abolished as arginine deprivation results in extensive oxidative damage (10). The case for drug discovery approaches to target arginine biosynthetic enzymes is further bolstered by work demonstrating that chemical inhibition of ArgJ with Pranlukast, a cysteinyl leukotriene receptor-1 antagonist use to treat asthmatic exacerbations, cleared a chronic *M. tuberculosis* infection in BALB/c mice (14). The arginine biosynthesis pathway consists of eight different enzymes (Figure 1A) all considered to be essential for *M. tuberculosis* growth *in vitro* (15). Except *argA* which encodes the first enzyme of the pathway, all other genes are present in a single operon that also includes the repressor *argR* (Figure 1B).

**Figure 1:**
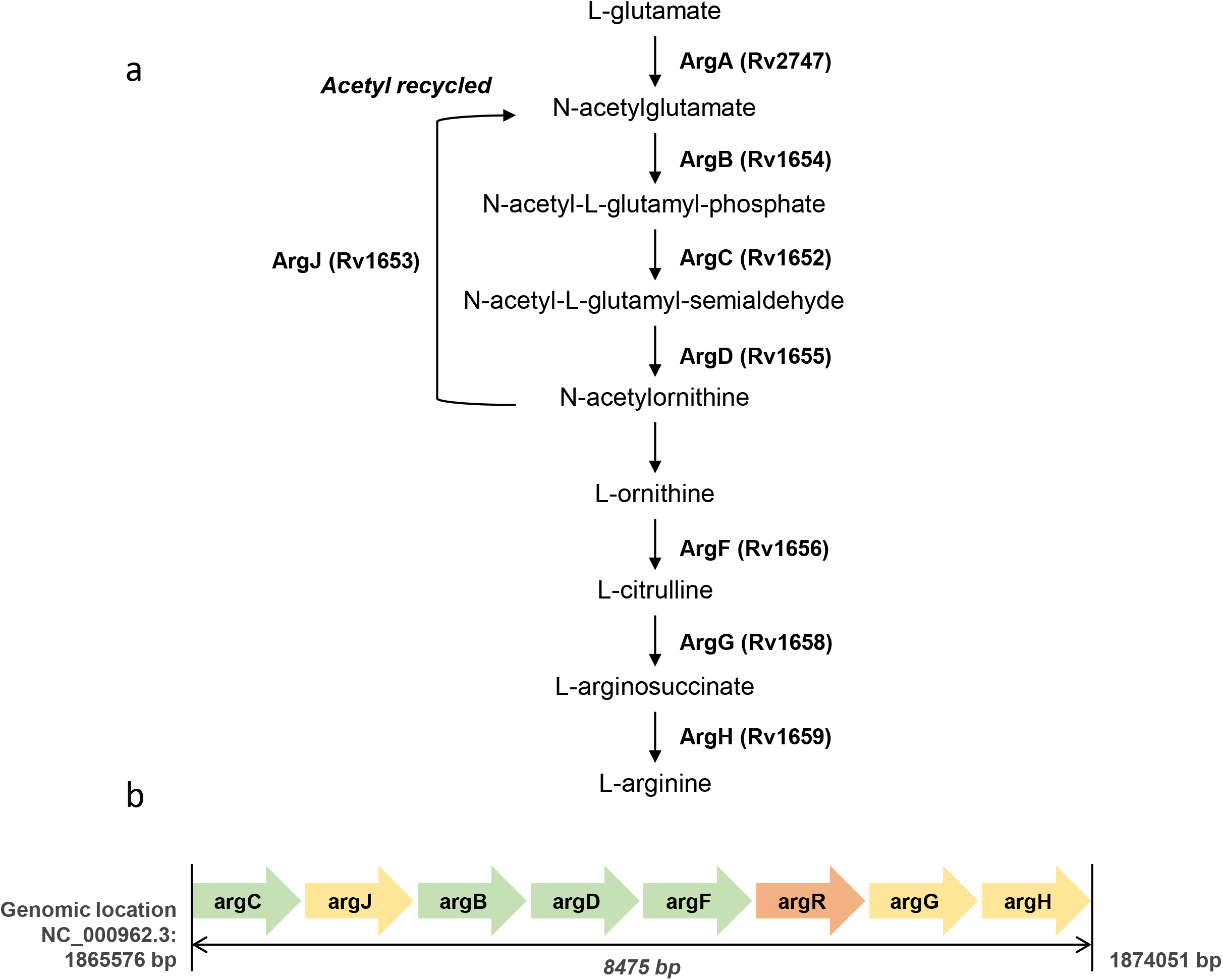
The L-arginine biosynthesis pathway in *Mycobacterium tuberculosis* (a). *M. tuberculosis* L-arginine biosynthesis operon (b).

Fragment based drug discovery (FBDD) is now an established lead-generation strategy in both industry and academia, having yielded over 30 compounds in clinical trials, including approved cancer drugs like vemurafenib, Kisqali, Balversa and venetoclax (16). This approach consists of screening a library of small molecules (150-300 Da) against a target of interest using biophysical, biochemical and structural biology methods. The low complexity of fragments allows for efficient exploration of the chemical space of the target, often revealing unexpected binding sites in proteins. Although fragments often bind weakly, they tend to bind to hotspot regions of the target, forming well defined interactions that allow subsequent elaboration into larger, drug-like molecules (17, 18). Our group and a few others have pioneered using this approach against different mycobacterial species and different protein targets with varying degrees of success (18–26).

Using this approach, we have screened four enzymes of the arginine biosynthesis pathway not yet explored drug discovery programmes: ArgB, ArgC, ArgD and ArgF. Herein we report the structures of the four enzymes in complex with fragments hits, including a novel allosteric site of ArgB and allosteric inhibitors of this enzyme. Importantly, this work also assesses the potential of these enzymes as candidates of future drug discovery programmes.

## 2. Materials and Methods

### 2.1. Molecular Cloning

The *argB* gene was amplified from chromosomal DNA of *M. tuberculosis* H37Rv strain obtained from ATCC (ATCC25618D-2) while the ORFs of *argC*, *argD* and *argF* were purchased as *E. coli* codon-optimised synthetic gene strings through the ThermoFisher GeneArt Gene Synthesis service. The *argB* gene was cloned into pHAT4 (27) using NcoI and XhoI sites. The gene strings of *argC*, *argD* and *argF* were cloned into a pET28a vector (modified to include an N-terminal 6xhis SUMO) (28) using BamHI and HindIII restriction sites. All constructs were confirmed by sequencing.

### 2.2. Protein expression and purification

250 mL of autoclaved 2xYT broth (Formedium) prepared in distilled water, containing 100 μg/mL ampicillin for pHAT4:*argB* or 30 μg/mL kanamycin for pET28a:*argC*/*argD*/*argF*, was inoculated with *E. coli* BL21(DE3) containing the respective expression construct, and incubated at 37 °C with 220 rpm shaking overnight. This primary culture was used the following day to inoculate 6 flasks containing 1 L 2xYT broth and the appropriate antibiotic, and the inoculated flasks were incubated under similar conditions until the OD_600nm_ reached 0.8-1. Overexpression was induced by the addition of 0.5 mM isopropyl β-D-1-thiogalactopyranoside (IPTG). Thereafter, the flasks were incubated at 20 °C with 220 rpm shaking overnight.

Cells were harvested by centrifugation at 4200 rpm, 4 °C for 20 minutes in a Beckman Coulter ultracentrifuge. The cell pellets were re-suspended in 50 mL of Buffer A (Table S1), also containing 1 tablet of cOmplete™, Mini, EDTA-free Protease Inhibitor Cocktail (Roche, Merck), DNase I (Sigma-Aldrich) and 5 mM MgCl_2_. The cells were lysed by ultrasonication for ~ 8 minutes (pulse on for 20 secs, pulse off for 30 secs, 55% amplitude), the suspensions were kept in an ice bath throughout. The cell lysates were clarified by centrifugation (27000 *g*, 4 °C for 40 minutes), and the supernatants were syringe-filtered (0.45 μm membrane) to remove any cell debris.

The filtered lysates were subjected to IMAC using a His-Trap 5 mL Nickel column (GE Healthcare Life Sciences) on an ÄKTA Pure system (GE Healthcare Life Sciences), equilibrated with buffer A (Table S1).

Isocratic elution was performed using buffer B (Table S1). Proteins were dialysed in buffer C at 4 °C and tags were cleaved overnight by adding TEV protease for ArgB or Ulp1 protease, for ArgC, ArgD and ArgF, both at 1:100 ratio.

The dialysed proteins were concentrated to a <5 mL volume using a 30 kDa MWCO Vivaspin 20 centrifugal concentrator (Sartorius) at 5000 *g*, 4 °C and injected onto a HiLoad 16/600 Superdex 200 gel filtration column (GE Life Sciences) equilibrated with buffer C (Table S1). Elution fractions corresponding to the peak of interest in the chromatogram were pooled and fraction purity was assessed by SDS-PAGE. The purest fractions of ArgB and ArgF were pooled and concentrated to 20 mg.ml^−1^. Pooled fractions of ArgC were further dialysed into the final storage buffer (5 mM Tris-HCl pH 7.4, 50 mM NaCl) overnight at 4 °C, rescued the next day and concentrated to 6.5 mg/mL. ArgD fractions were pooled and aqueous pyridoxal-5’-phosphate (PLP, Sigma-Aldrich) was added (2 mM final PLP concentration). Overnight dialysis into the storage buffer (50 mM Tris-HCl pH 7, 100 mM NaCl) was carried out at 4 °C to remove excess PLP. The PLP-saturated protein was rescued the next day and concentrated to 14 mg/mL. All proteins were flash frozen in liquid N_2_ and stored at −80 °C.

### 2.3. Differential Scanning Fluorimetry

Fragment screening was carried out in a 96-well PCR plate using a CFX Connect Real-time PCR Detection System (Bio-Rad) for DSF. For ArgB, each 25 μL reaction mixture contained 10 μM ArgB, 100 mM HEPES (pH 7.5), 200 mM NaCl, 5x SYPRO Orange, 5% DMSO (v/v), and fragments at 5 mM. For ArgF each 25 μL reaction mixture contained 5 μM ArgF, 25 mM Tris-HCl (pH 8.0), 150 mM NaCl, 5x SYPRO Orange dye, 5 mM fragments and 5% DMSO (v/v). For ArgC and ArgD, the 25 μL reaction volume consisted of the following: 2.5 μM ArgC/5 μM ArgD, 100 mM sodium phosphate pH 7, 200 mM NaCl, 5x SYPRO Orange, and 5 mM fragments (960 fragment library). The protocol implemented increased temperature by 0.5 °C after every 30 seconds, going from 25 °C to 95 °C and measuring SYPRO Orange fluorescence for each temperature cycle. The melt curve RFU (relative fluorescence units) and derivative -d(RFU)/dT values were analysed and plotted using a macros-enabled Excel Workbook: the minima of the melt curves were recorded as the melting temperature (T_m_) of the enzymes in the presence of each fragment. The T_m_ of the reference control (protein in the presence of DMSO) was subtracted from all the readings to calculate ΔT_m_.

### 2.4. Surface plasmon resonance

Low molecular weight (LMW) screening with the DSF fragment hits was carried out using the T200 Biacore instrument (GE Healthcare Life Sciences). A series S CM5 sensor chip (GE Healthcare Life Sciences) was used for the immobilisation of ArgC on the carboxymethylated dextran matrix through amine coupling. A 25 μg/mL ArgC dilution was prepared in the optimal coupling buffer (Sodium acetate pH 5), and immobilisation was performed by manual instructions. The immobilised ArgC was tested using dilution series (19 μM to 2.5 mM) of NADP^+^ and NADPH. 50 mM fragment DMSO stocks were used to prepare 1 mM dilutions in the SPR buffer consisting of 10 mM sodium phosphate pH 7.0, 150 mM NaCl and 2% (v/v) DMSO. Each NADP^+^/NADPH and fragment dilution was injected once at a flow rate of 30 μL/min for a contact time of 30 seconds, SPR Running buffer, consisting of 10 mM sodium phosphate pH 7.0 and 150 mM NaCl, was passed for 320 seconds at the same flow rate, and 50% DMSO (diluted in SPR running buffer) was injected at the end of the cycle to remove undissociated analyte. Solvent correction was carried out to account for DMSO mismatch between the analyte dilutions and the SPR running buffer.

A 30 μg/mL ArgD dilution was prepared in sodium acetate pH 4 buffer for immobilisation. Following the ethanolamine neutralisation step, 1 mM PLP (prepared in the SPR buffer) was injected for a contact time of 420 seconds to ensure saturation of PLP-binding sites. The immobilised ArgD was tested using dilution series (19 μM to 2.5 mM) of L-glutamate, N-acetylornithine and L-ornithine. Screening was also carried out against PLP-unsaturated ArgD.

### 2.5. Ligand-observed NMR

All NMR experiments were carried out at 298 K using a Bruker Avance 600 MHz spectrometer with a Triple Resonance Inverse (TCI) Automatic Tuning and Matching (ATM) cryoprobe. T2 relaxation-filtered one-dimensional NMR spectroscopy experiments incorporated a CPMG67 spin-lock time of 200 ms before the acquisition period. Samples (600 μL) containing 2 mM fragment in the absence and presence of 10 μM ArgB were prepared in buffer containing 20 mM potassium phosphate at pH 7.4 and 50 mM NaCl. Additionally, 2% v/v d6-DMSO was present in all samples for fragment solubilisation and field-frequency locking. Displacement experiments were carried out in the same manner by adding 1 mM each of ATP, and N-acetyl-L-glutamic acid or L-arginine to the samples containing 2 mM fragment and 10 μM ArgB. The samples were loaded into 5 mm NMR tubes (Wilmad, 526-PP) for measurement, and the resulting spectra were analysed using TopSpin v. 3.5 (Bruker).

### 2.6. Crystallisation of the apoenzymes

Crystallisation screening and optimisation for all the enzymes was performed at 18 °C with the sitting drop vapour diffusion method using a Mosquito robot (TTP-Labtech) to setup the crystallisation experiments. For apo ArgB, 300 nL of pure protein at 10 mg.ml^−1^ was mixed with an equal volume of reservoir solution and equilibrated against 85 μl of the reservoir solution. The selected condition was obtained in Wizards Classic 1&2 crystallisation screen (Rigaku), well G5 (1260 mM ammonium sulphate, 100 mM CHES pH 9.5 and 200 mM NaCl). Crystals appeared after 2 days in this condition and diffracted up to 1.8 Å resolution. For ArgF, 200 nL of pure protein at 20 mg.mL^−1^ was mixed with an equal volume of reservoir solution. An initial crystallization condition was identified I the Wizards Classic 3&4 crystallisation screen (Rigaku), well F2 (40% PEG400, 100 mM Tris-HCl pH 7.5 and 200 mM Lithium sulphate). However, crystals obtained in this condition contained two lattices with different orientations and no structural solution could be found despite good quality diffraction. These crystals were ground to produce seeds and a new crystallisation screening was performed using 200 nL of ArgF at 20 mg.mL^−1^ mixed with equal volume of reservoir solution and 50 nL of seed solution. After several rounds of optimisation, the final crystallisation condition consisted of 150 mM ammonium dihydrogen phosphate and 10 mM praseodymium acetate. Crystals appeared after a 2 days and diffracted up to 1.8 Å resolution.

A previously reported crystallisation condition for ArgC (29) was reproduced with some modifications, but the crystals were found to not be suitable for fragment soaking experiments. A new crystallisation screen was therefore performed using 200 nL of pure ArgC at 6.5 mg.ml^−1^ mixed with an equal volume of reservoir solution. Well A8 from the BCS screen (Molecular Dimensions) produced crystals diffracting to 1.54 Å. This condition consisted of 0.1 M phosphate/citrate buffer pH 5.5 and 20% PEG Smear High (PEG 6K, 8K, 10K) and was optimised to remove the cryoprotection step by adding 20% glycerol. A second condition with a neutral pH more amenable to soaking based on the previous condition was also optimised and consisted of 0.1 M Bis-Tris pH 7, 17% PEG Smear High, 70 mM phosphate/citrate pH 5.6, 20% glycerol.

For ArgD, 200 nL PLP-saturated enzyme 14 mg.ml^−1^ was mixed with an equal volume of reservoir solution. A condition was found in PEG Smear BCS screen (Molecular Dimensions) well F6 (0.1 M Bis-Tris Propane pH 8.5, 18% PEG Smear High (PEGs 6K, 8K, 10K), 0.2 M Ammonium nitrate). The final optimized conditions consisted of 0.1 M Bis-Tris Propane pH 8.5, 18% PEG Smear High, 0.2 M ammonium nitrate and 10 mM nickel chloride (additive).

### 2.7. Crystal soaking and co-crystallisation with natural ligands and fragment hits

To obtain ligand-bound structures, soaking was performed in the crystallisation conditions described above for each protein using the hanging drop vapour diffusion method.

For ArgB, 1.5 μL of protein storage buffer containing 20 mM of ligand was mixed with 1.5 μL of reservoir solution, and drops were left to equilibrate against 500 μL of reservoir solution for 3 days. Crystals were then transferred to the drops and incubated for 16 h. A cryogenic solution was prepared by adding ethylene glycol up to 27.5% v/v to the mother liquor. Crystals were briefly transferred to this solution, flash-frozen in liquid N_2_, and stored for data collection. To obtain an ArgB-NAG complex, co-crystallization with 2 mM NAG was performed instead. Crystals for ArgB-NAG complex were obtained in Wizard Classic 1&2 screen (Rigaku), solution B6, and were flash-frozen in liquid N_2_ after a brief soak in a solution containing mother liquor and 27.5% ethylene glycol.

ArgC crystals grown in pH 5.5 were first soaked in 1.5 μL drops containing the mother liquor and 5 mM NADP^+^ for 2 hours in hanging drops that were equilibrated against a reservoir of 500 μL. Thereafter, the crystals were transferred to drops containing the crystallisation condition and an SPR-validated fragments (20 mM, 10% DMSO), which were equilibrated against 500 μL of mother liquor also containing a corresponding percentage of DMSO overnight at 19 °C. ArgC crystals grown in pH 7 were soaked with 5 mM NADP^+^ only for 5-10 minutes due to the rapid development of cracks, and transferred to the fragment soaking drops for 5-10 minutes from where they were fished and frozen. ArgD crystals were soaked with fragments at a concentration of 50 mM overnight in otherwise the same manner as ArgC crystals grown in pH 5.5. A cryogenic solution was prepared by adding 30% ethylene glycol to the mother liquor. Crystals were briefly transferred to this solution and flash-frozen in liquid N_2_.

ArgF crystals were soaked in drops containing crystallization condition and 20 mM of ligand and equilibrated against 500 μL of reservoir solution for 16h. A cryogenic solution was prepared by adding 25% ethylene glycol to the mother liquor. Crystals were briefly transferred to this solution and flash-frozen in liquid nitrogen.

### 2.8. X-ray data collection and processing

X-ray diffraction data (single-wavelength anomalous diffraction) were collected on beamlines i02, i03, i04, i04-1 and i24 at the Diamond Light Source (DLS), UK and on id30B at The European Synchrotron Radiation Facility (ESRF). Diffraction data were processed and reduced with autoPROC from Global Phasing Limited (30) or Xia2 (31). The apo-form of ArgB was crystallized in the R3_2_ spacegroup with one protomer per asymmetric unit (ASU) and the ArgB:NAG complex in the P6_3_ spacegroup with two protomers per ASU. ArgF was crystallized in the P2_1_ spacegroup with 6 protomers in the ASU. ArgC was crystallized in C2 and P2_1_ spacegroups, with 2 and 4/8 protomers per ASU respectively. ArgD was crystallised in the P2_1_ space group as well but with 4 protomers in the ASU.

Initial phases were determined with PHASER (32) from PHENIX software package (33) using the *M. tuberculosis* ArgB structure (PDB: 2AP9), *M. tuberculosis* ArgF structure (PDB: 2P2G), *M. tuberculosis* ArgC structure (PDB: 2I3G) and *E. coli* Succinyl-ornithine transaminase (AstC, 42% sequence identity, PDB: 4ADB) as a search models, respectively for ArgB, ArgF, ArgC and ArgD. Model building was done with Coot (34), and refinement was performed in PHENIX (33, 35) for ArgB, ArgF, and ArgC. For ArgD, after the initial molecular replacement solution and a single cycle of refinement, PHENIX AutoBuild was used to generate a model for *M. tuberculosis* ArgD that was then refined with Coot and PHENIX. Structure validation was performed using Coot and PHENIX tools (33, 34). All figures were prepared with PyMOL (The PyMOL Molecular Graphics System, Version 2.0 Schrodinger, LLC).

### 2.9. Isothermal titration calorimetry

Binding interaction between ArgB or ArgF and ligands was characterised at 25 °C, using a Microcal ITC200 titration calorimeter (Microcal). An ArgB concentration between 75-150 μM was used for all titrations. Ligands (0.75-2 mM) were injected in 1.5 μl aliquots with 150 s spacing between injections for compound 1 and 110 s for all the others. For compound 2, L-canavanine and L-arginine two titrations were concatenated. Titration data was recorded in 25 mM HEPES pH 7.4 with 200 mM NaCl. Data were analysed by fitting a simple single-site model using Origin software (Microcal) (NMR711 and NMR446) or a six-site sequential binding model (ArgB and L-canavanine).

ArgF was dialysed in 50 mM HEPES pH 8.0, 200 mM NaCl before it was loaded into the calorimetry cell at concentrations of 75-100 μM with the addition of 1 mM DTT. Ligand solutions at concentrations of 1 mM were dissolved in the same buffer and typically injected at between 0.5 μL and 2 μL at 150 second intervals with stirring at 750 rpm. Buffer-ligand titrations were carried out as reference runs and subtracted from the protein-ligand titration to remove the heat of dilution. Data were analysed by fitting a simple single-site model using Origin software (Microcal).

### 2.10. Enzymatic assays

ArgB activity was assessed by a colorimetric assay that followed the release of ADP by measuring the oxidation of NADH at 340 nm, for 30 min, in the presence of pyruvate kinase and lactate dehydrogenase in a PHERAstar plate-reader (BMG-Labtech). The enzymatic reactions (200 μl) were performed at 25 °C and contained 50 mM Tris pH 7.5, 200 mM NaCl, 50 mM KCl, 10 mM MgCl_2_, 0.3 mM NADH, 2.5 mM phosphoenolpyruvate, 0.3 mM ATP, 1.25 mM N-acetyl-L-glutamate (NAG), 10% DMSO (v/v), 4 units of pyruvate kinase/lactate dehydrogenase, 0.5 μM ArgB, and varying concentration of inhibitors. Inhibitors were also individually screened against the coupled enzymes to eliminate any compounds interfering with the other assay components. Competition assays were performed in the same conditions using 0.3 mM of ATP or NAG and varying the other substrate concentration.

To synthesise the ArgC substrate a reaction mixture containing 3 μM ArgB, 1 mM NAG and 1 mM ATP in 50 mM Tris-HCl pH 7.5, 100 mM NaCl and 40 mM MgCl_2_ was made and the reaction was allowed to proceed for 1-1.5 hours at room temperature. Thereafter, 100 μL of the ArgC reaction mixture consisting of 50 mM Tris-HCl pH 7.5, 100 mM NaCl and 0.6 mM NADPH (concentration in 200 μL), was added to each well together with 100 μL of the ArgB reaction mixture and followed for 30 min at 35 °C by measuring the oxidation of NADPH at 340 nm. Controls with only NADPH, 2.5% DMSO and no ArgC, as well as only NADPH, 4 mM fragment and no ArgC were prepared. Baseline ArgC activity was assayed with and without DMSO, the effect of 2.5% and 5% DMSO on enzymatic activity was also tested. To test the inhibitory effect of fragment binders identified from the crystallographic screening, 6 fragment concentrations (125 μM, 250 μM, 500 μM, 1 mM, 2 mM and 4 mM,) were added to the reaction mixtures from suitable DMSO stocks such that the final DMSO concentration was 2.5%. All conditions were prepared in triplicates. 3.5 μM ArgC was added just before measurements were started.

To assess the activity of ArgD an end point assay that follows the reverse reaction of the enzyme was used in the presence and absence of fragments. Reactions (100 μL) were set up containing the following: 100 mM Tris-HCl pH 8, 3 mM N-acetylornithine (NAO), 3.4 mM α-KG, 20 μM PLP and 4 μM enzyme. Duplicate reaction mixtures were prepared for each time point and 8 time points (0, 5, 10, 15, 20, 30, 45, 60 minutes) were tested in total. Two controls were prepared: one with all reaction components except the enzyme, and the other with all reaction components except NAO (NAO was added after HCl treatment). The reactions tubes were allowed to equilibrate in a heating block set at 37 °C for two minutes, and the enzyme was added to initiate the reaction. After the stipulated incubation times, contents of the reaction tubes were quickly transferred to 1.5 mL microcentrifuge tubes containing 60 μL of 6 N HCl to stop the reactions. These tubes were then kept in a heating block set at 95 °C for 30 minutes, after which they were cooled to 25 °C in a water bath. 200 μL of 3.6 M sodium acetate was added to each tube (final concentration of 1.8 M), along with 40 μL of 30 mM 2-aminobenzaldehyde (final concentration of 3 mM in a total volume of 400 μL). A yellow colouration started developing as soon as the latter was added, the contents were vortexed and the tubes were incubated at 25 °C for 15 minutes. 200 μL of each reaction mixture was transferred into wells of a 96-well flat bottom UV transparent microplate, and absorbance at 440 nm was measured using the PHERAstar plate reader. All experiments were performed at least in triplicate in a PHERAstar plate-reader (BMG-Labtech) and the control without ArgD was used for blank subtraction. Data were analysed with GraphPad Prism (Graphpad Software). All reagents were obtained from Sigma-Aldrich.

### 2.11. *M. tuberculosis* culture condition and minimum inhibitory concentration (MIC) determination

Mutant strain Δ*argB* and its complemented strain Δ*argB*-c were generated before as mentioned (10). All the strains, wild type *M. tuberculosis* H37Rv and mutants Δ*argB* as well as Δ*argB*-c were grown at 37 °C to mid-log phase in Middlebrook 7H9 medium supplemented with 10% oleic acid-albumin-dextrose-catalase (OADC), 0.5% glycerol, and 0.05% Tyloxapol supplemented with arginine (1mM) washed and suspended in 7H9 media ± arginine (1mM). For MIC, the cultures were diluted to 1/500 in ± arginine supplemented media. Serial two-fold dilutions of each drug were prepared directly in a sterile 96-well plate using 0.1 ml of media with the appropriate supplement in the presence or absence of 1mM arginine. Same media with only vehicle (no drug) was used as a control. PBS (0.2 ml) was added to all the perimeter wells. The diluted *M. tuberculosis* strains in ± arginine supplemented media (0.1 ml) were added to each well, and the plate was incubated at 37°C for 7 days. Cell growth was measured by optical density at 600 nm. An aqueous solution of resazurin (0.2 mg/ml; 0.03 ml) was added to each well, and the plate was further incubated for up to two days at 37°C. The MIC was determined as the lowest concentration at which the change of colour from blue (resazurin) to pink (resorufin) did not occur.

## 3. Results

### 3.1. DSF fragment screening

In the first stage of the screening, differential scanning fluorimetry was used to screen an in-house library of 960 rule-of-three compliant fragments against four enzymes of the arginine biosynthesis pathway ArgB, ArgC, ArgD and ArgF. In the case of ArgB, ArgC and ArgF the screening was performed against the apoenzymes while for ArgD it was done with the PLP-bound form. Several known ligands (substrates, products, allosteric regulators and co-factors) were used as positive controls for each of the enzymes. In all cases, a fragment was considered a hit when the shift in melting temperature was greater than five times the standard deviation.

#### 3.1.1. ArgB

In the conditions used in the assay, ArgB with 5% DMSO displayed a melting temperature of 48 °C. The addition of 1 mM ATP, N-acetyl-glutamate and L-arginine showed positive melting shifts of 2.0, 11.8 and 8.6 °C respectively. Of the 960 compounds, a total of 63 (≈6.6%) showed a thermal shift greater than five standard deviations (≥ 1.25 °C) at 5 mM and were considered hits. Out of those, 14 showed a large stabilization of ArgB with a thermal shift greater than 5 °C (Table 1) and were selected for further validation.

**Table 1:**
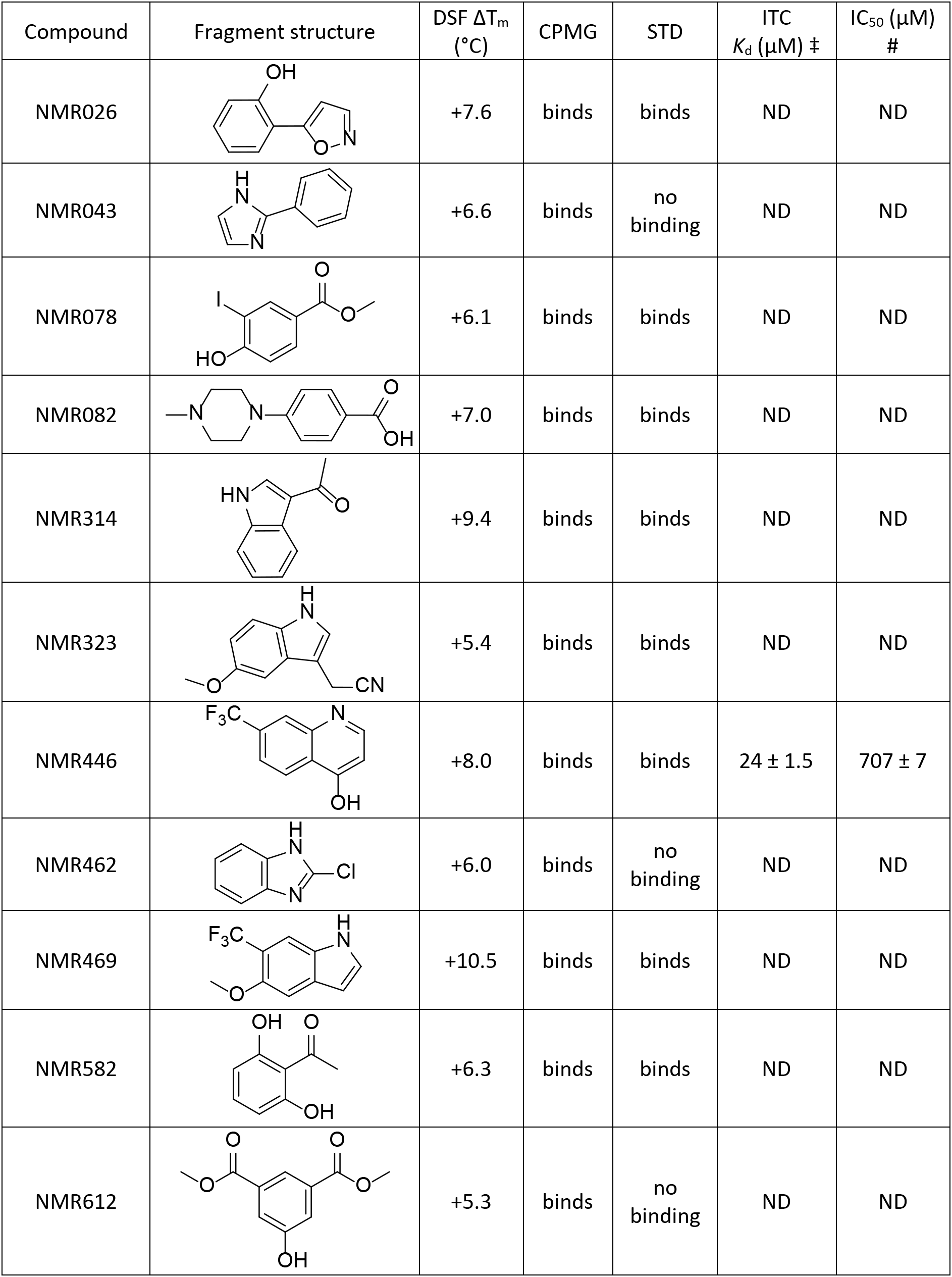
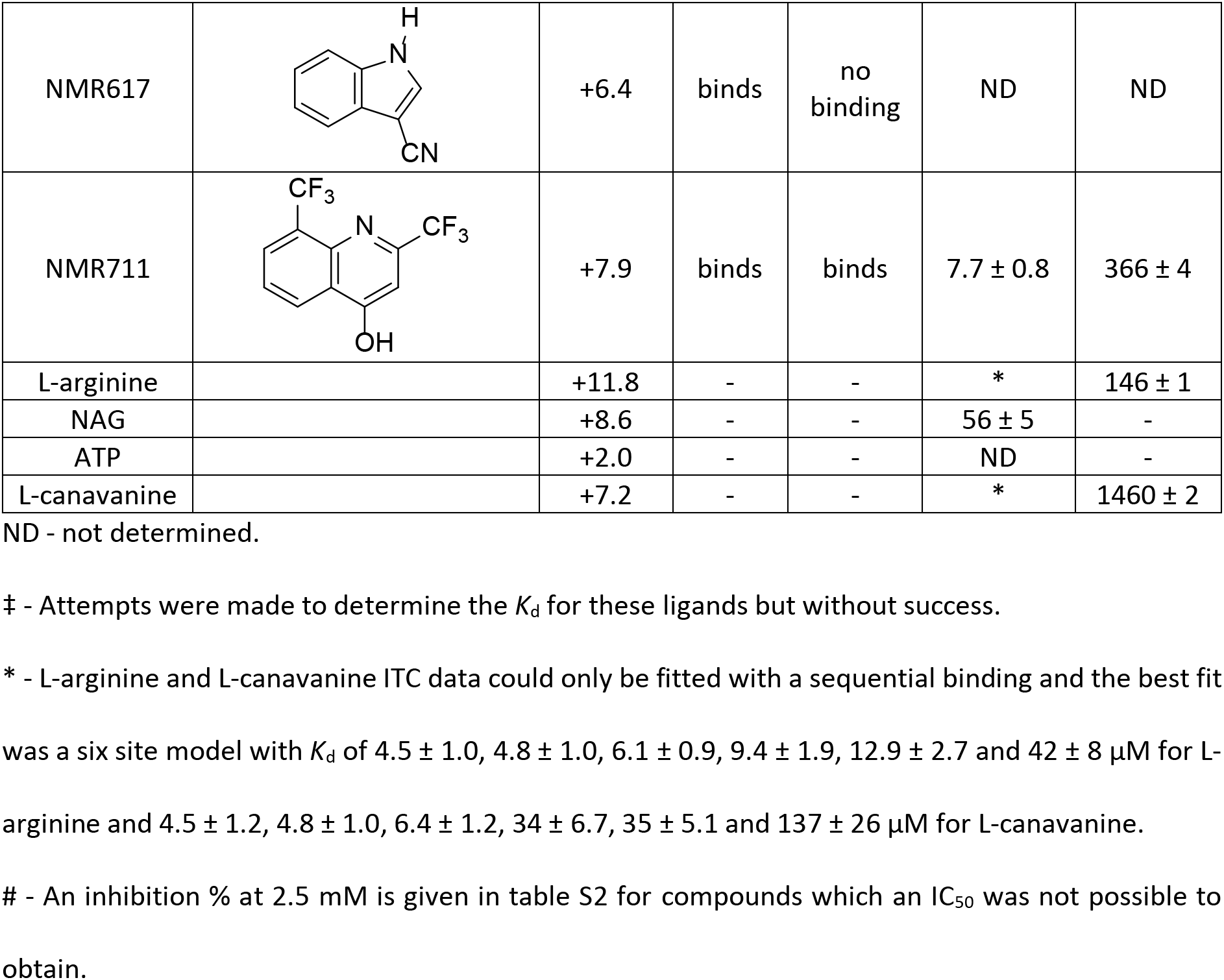
ArgB validated fragment hits.

| Compound | Fragment structure | DSF $\Delta T_m$<br>(°C) | CPMG | STD | ITC<br>$K_d$ (μM) ‡ | IC<sub>50</sub> (μM)<br># |
| --- | --- | --- | --- | --- | --- | --- |
| NMR026 |  | +7.6 | binds | binds | ND | ND |
| NMR043 |  | +6.6 | binds | no binding | ND | ND |
| NMR078 |  | +6.1 | binds | binds | ND | ND |
| NMR082 |  | +7.0 | binds | binds | ND | ND |
| NMR314 |  | +9.4 | binds | binds | ND | ND |
| NMR323 |  | +5.4 | binds | binds | ND | ND |
| NMR446 | | +8.0 | binds | binds | 24 ± 1.5 | 707 ± 7 |
| NMR462 |  | +6.0 | binds | no binding | ND | ND |
| NMR469 |  | +10.5 | binds | binds | ND | ND |
| NMR582 |  | +6.3 | binds | binds | ND | ND |
| NMR612 |  | +5.3 | binds | no binding | ND | ND |
| NMR617 |  | +6.4 | binds | no binding | ND | ND |
| NMR711 | | +7.9 | binds | binds | $7.7 \pm 0.8$ | $366 \pm 4$ |
| L-arginine | | +11.8 | - | - | * | $146 \pm 1$ |
| NAG | | +8.6 | - | - | $56 \pm 5$ | - |
| ATP |  | +2.0 | - | - | ND | - |
| L-canavanine | | +7.2 | - | - | * | $1460 \pm 2$ |

ND - not determined.
‡ - Attempts were made to determine the $K_d$ for these ligands but without success.
\* - L-arginine and L-canavanine ITC data could only be fitted with a sequential binding and the best fit was a six site model with $K_d$ of $4.5 \pm 1.0$ , $4.8 \pm 1.0$ , $6.1 \pm 0.9$ , $9.4 \pm 1.9$ , $12.9 \pm 2.7$ and $42 \pm 8 \mu\text{M}$ for L-arginine and $4.5 \pm 1.2$ , $4.8 \pm 1.0$ , $6.4 \pm 1.2$ , $34 \pm 6.7$ , $35 \pm 5.1$ and $137 \pm 26 \mu\text{M}$ for L-canavanine.

#### 3.1.2. ArgC

The apoenzyme ArgC with 5% DMSO was found to have a melting temperature of 67.7 °C. While 1 mM NADP^+^ gave a small average positive thermal shift of ~0.6 °C and 1 mM NADPH produced an average negative thermal shift of −0.9 °C across all DSF runs performed. 81 out of the 960 fragments screened (≈8.5% of the library) at a concentration of 5 mM gave a positive thermal shift greater than five standard deviations (≥ 2.9 °C). An orthogonal biophysical technique, Surface Plasmon Resonance (SPR), was also employed to corroborate these hits (Table 2).

**Table 2:**
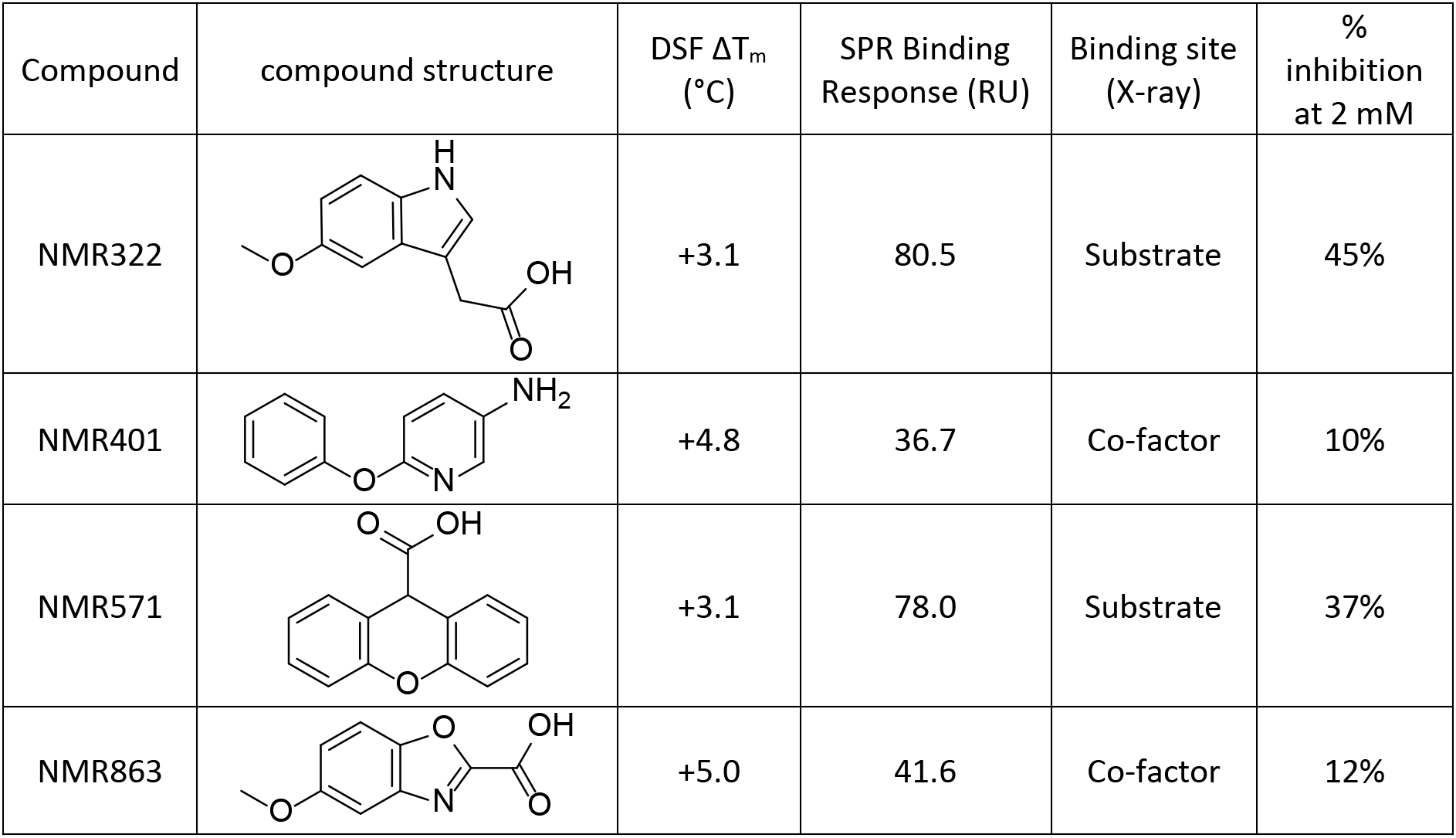
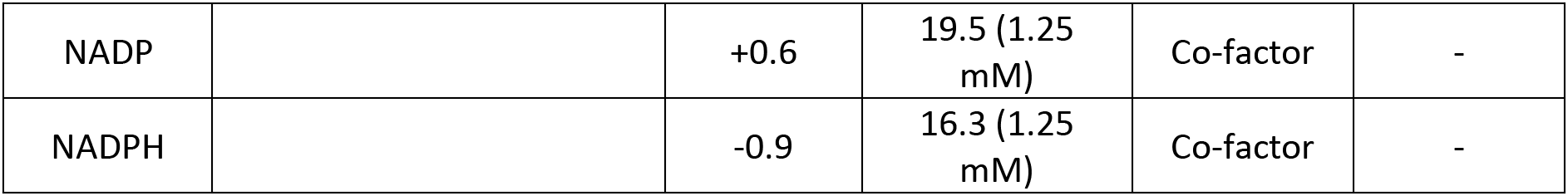
ArgC validated fragment hits.

| Compound | compound structure | DSF $\Delta T_m$ ($^{\circ}\text{C}$) | SPR Binding Response (RU) | Binding site (X-ray) | % inhibition at 2 mM |
| --- | --- | --- | --- | --- | --- |
| NMR322 |  | +3.1 | 80.5 | Substrate | 45% |
| NMR401 |  | +4.8 | 36.7 | Co-factor | 10% |
| NMR571 |  | +3.1 | 78.0 | Substrate | 37% |
| NMR863 |  | +5.0 | 41.6 | Co-factor | 12% |
| NADP |  | +0.6 | 19.5 (1.25 mM) | Co-factor | - |
| NADPH |  | -0.9 | 16.3 (1.25 mM) | Co-factor | - |

#### 3.1.3. ArgD

ArgD contains the prosthetic group PLP that does not leave the active site of the enzyme. Therefore, all the PLP sites must remain occupied during screening experiments to represent the native state of the target. In an attempt to saturate all of the ArgD PLP sites, PLP was added to ArgD during the purification of the protein. Further confirmation was required to assess if most of the sites were now saturated. To examine this, the melting profiles of unsaturated ArgD, saturated ArgD and the two forms in the presence 1 mM PLP were assessed. Unsaturated ArgD exhibited a melting profile with 2 peaks, a large and broad peak with melting temperature of 64.5 °C, and a small peak with a melting temperature of 77 °C (Figure S1). The addition of 1 mM PLP changed the melting profile to a single peak with a melting temperature to 77.5 °C (Figure S1). ArgD that had PLP added during purification showed a melting temperature of 77.5 °C (Figure S1). The addition of 1 mM PLP had now a very minor stabilizing effect shifting the melting temperature to 79 °C (Figure S1) and confirming that most PLP sites were saturated. However, the protein in this state was insensitive to fragment binding with maximum thermal shifts of 0.5 °C being observed. We therefore tested the potential of using the unsaturated protein for the screening. As mentioned above, PLP-unsaturated ArgD exhibited a melting profile with 2 peaks, a large and broad peak with melting temperature of 64.5 °C (Figure S1), likely a mixture of two different populations in which none or one of the two protomers contain PLP, and a small peak with a melting temperature of 77 °C (Figure S1) which most likely corresponded to a PLP-saturated population. The fact that the addition of 1 mM PLP shifts the melting profile to a single peak with a melting temperature of 77.5 °C corroborated this hypothesis.

Three different types of response in the melting profile to the presence of fragments were observed while screening the PLP-unsaturated ArgD. Most fragments showed either no effect on the melting profile or a decrease in the melting temperature of the large peak or of both peaks and were discarded (Figure S1). A second set caused a change in the melting profile with a large increase in the intensity of the highest temperature peak (Figure S1) suggesting that the fragment was preferentially binding to the PLP binding site. The third set, had fragments that led to an increase in melting temperature inferior to 4 °C but maintained the melting profile of the PLP-unsaturated ArgD control (Figure S1). Fragments of this set could either be binding to the PLP site but not stabilizing the protein sufficiently to show a clear change in the melting profile, or could be binding elsewhere on the protein both in the presence or absence of PLP. This set, represented by 47 fragments (≈4.9% of the library) giving melting shifts of at least five standard deviations (≥ 2 °C) were therefore selected for SPR validation (Table 3).

**Table 3:**
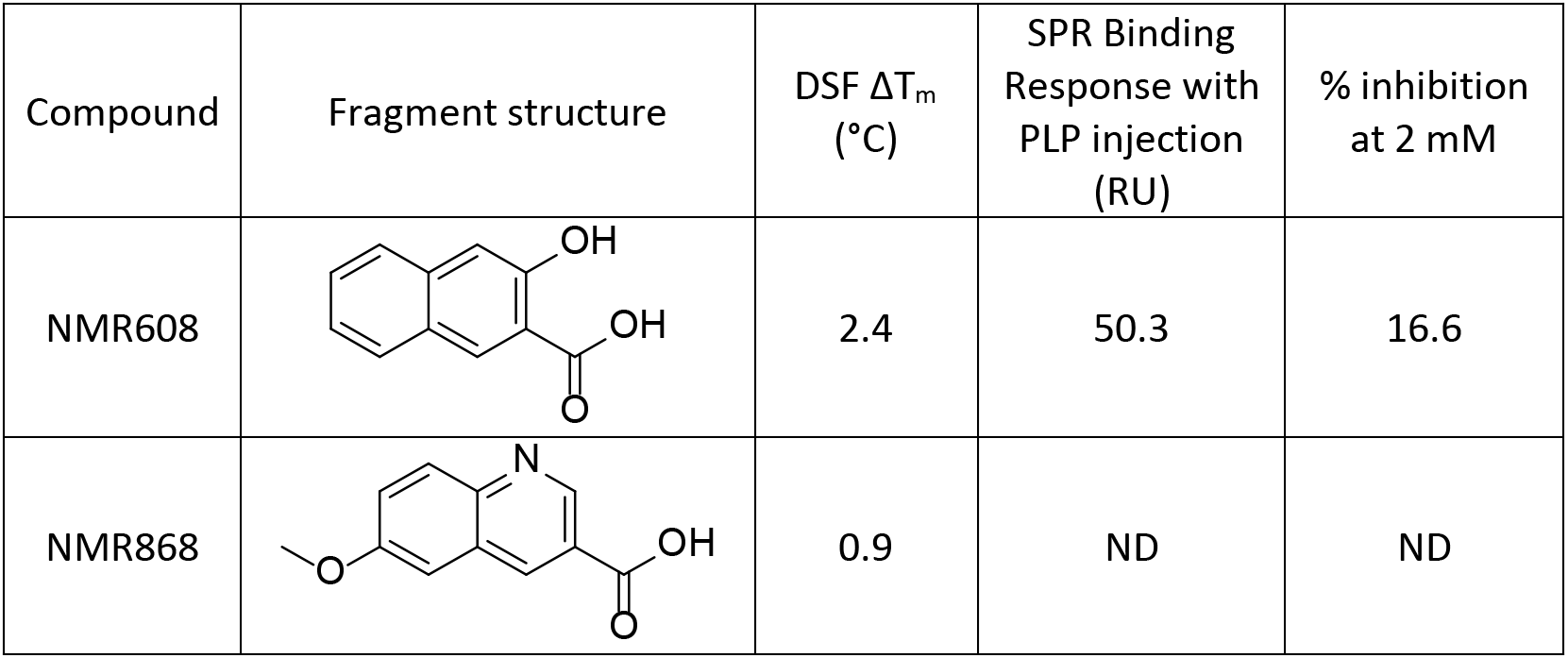
ArgD validated fragment hits.

| Compound | Fragment structure | DSF $\Delta T_m$ (°C) | SPR Binding Response with PLP injection (RU) | % inhibition at 2 mM |
| --- | --- | --- | --- | --- |
| NMR608 |  | 2.4 | 50.3 | 16.6 |
| NMR868 |  | 0.9 | ND | ND |

#### 3.1.4. ArgF

ArgF displayed a melting temperature of 65.5 °C with 1 mM L-ornithine and L-citrulline showing a positive melting shift of 1.5 and 1 °C respectively. Of the 960 fragments, a total of 105 (≈10.9%) showed a thermal shift greater than five times standard deviations (≥ 1.0 °C) at 5 mM and were considered hits. Out of these 16 displayed a melting shift greater than 3 °C and were selected for further validation by X-ray. Due to the large number of hits for this protein, greater than 10% of the whole library a clustering analysis of the fragment hits was performed. Centroids for each identified cluster and the representative displaying the highest melting shift were also selected for X-ray validation (Table 4).

**Table 4:**
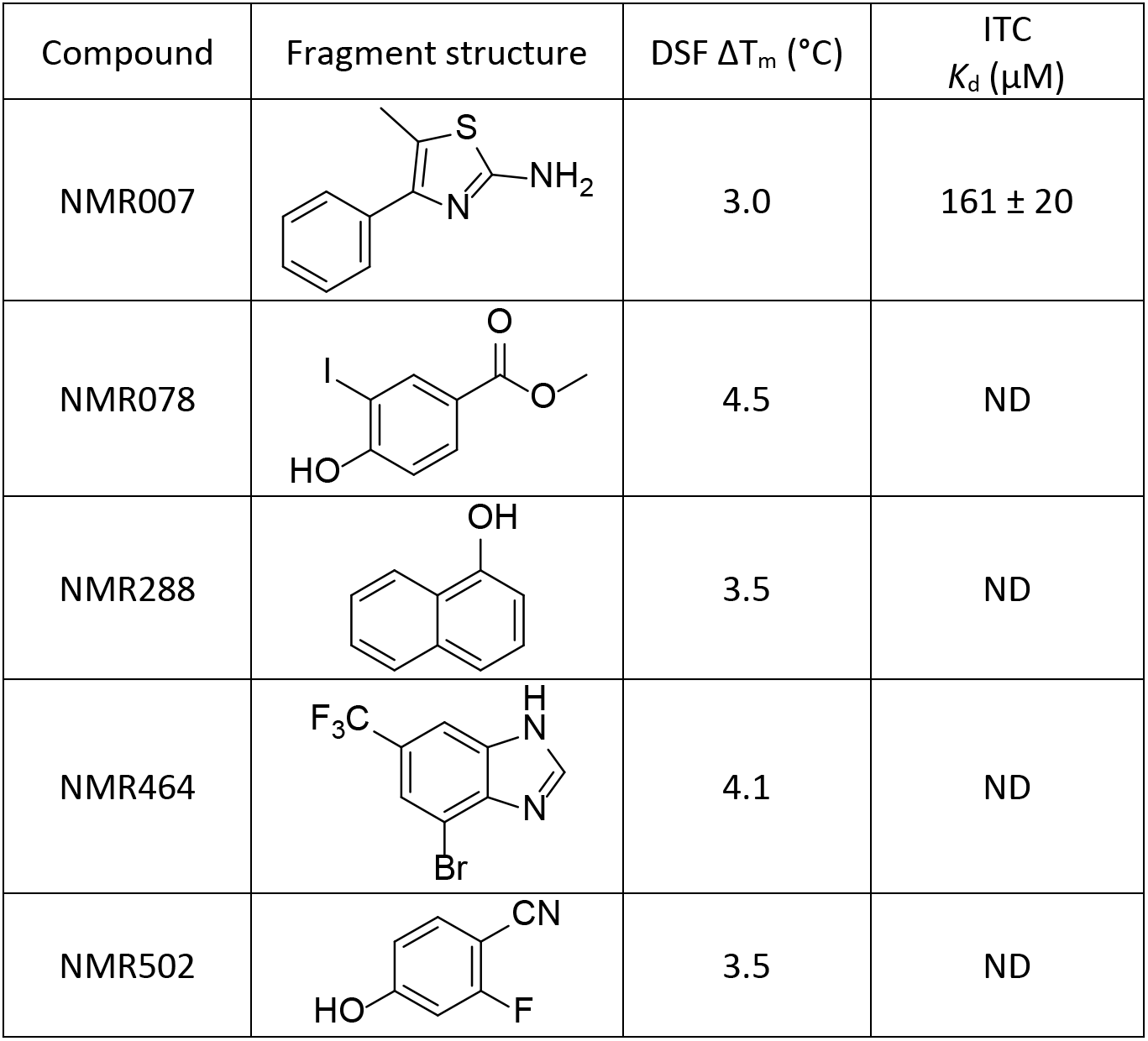
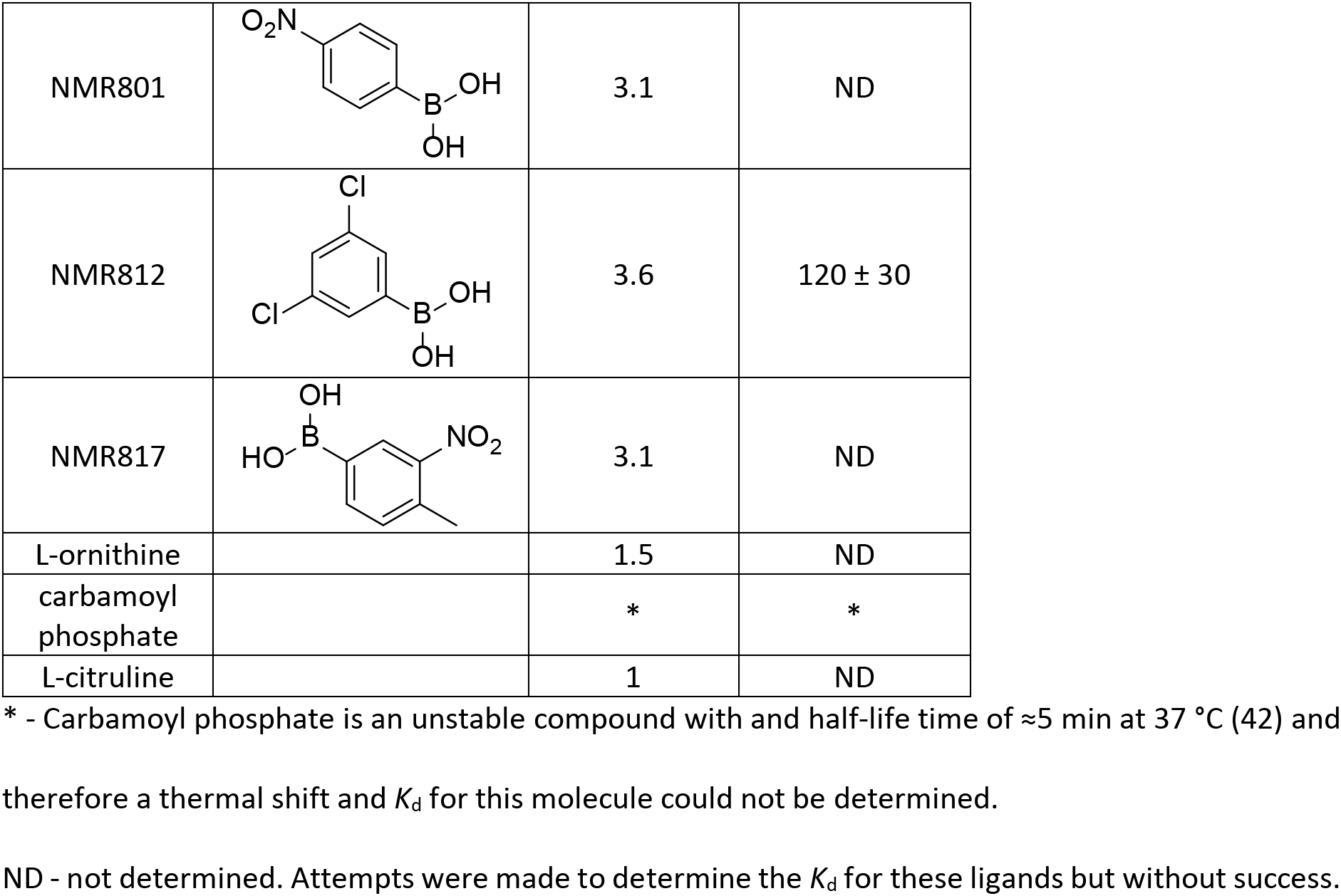
ArgF validated fragment hits.

| Compound | Fragment structure | DSF $\Delta T_m$ (°C) | ITC $K_d$ (μM) |
| --- | --- | --- | --- |
| NMR007 |  | 3.0 | 161 ± 20 |
| NMR078 |  | 4.5 | ND |
| NMR288 |  | 3.5 | ND |
| NMR464 |  | 4.1 | ND |
| NMR502 |  | 3.5 | ND |
| NMR801 |  | 3.1 | ND |
| NMR812 |  | 3.6 | 120 ± 30 |
| NMR817 |  | 3.1 | ND |
| L-ornithine |  | 1.5 | ND |
| carbamoyl phosphate |  | * | * |
| L-citruline |  | 1 | ND |

\* - Carbamoyl phosphate is an unstable compound with a half-life time of $\approx 5$ min at 37 °C (42) and therefore a thermal shift and $K_d$ for this molecule could not be determined.
ND - not determined. Attempts were made to determine the $K_d$ for these ligands but without success.

### 3.2. Validation of DSF hits using a secondary screening technique

Three different strategies were employed to validate the hits obtained with DSF. For ArgF the fragment hits were taken directly for X-ray crystallographic validation, for ArgB ligand-based NMR was employed and for ArgC and ArgD SPR was performed to validate the DSF hits.

#### 3.2.1. ArgB

To validate the ArgB hits obtained by DSF two ligand based NMR methods, Carr-Purcell-Meiboom-Gill (CPMG) and STD (saturated transfer difference), were used (36, 37) and fragments that were validated by at least one method were considered as confirmed hits. CPMG experiments validated 15 out of the 16 fragments while CPMG validated 11 out of 16. Only one fragment was not validated by both methods and thus was not taken forward.

#### 3.2.2. ArgC and ArgD

ArgC and PLP saturated ArgD were immobilised on an activated carboxymethylated dextran surface via amine coupling and final immobilisation response achieved was ≈16000 RU for ArgC and ≈7500 RU for ArgD (1000 RU roughly corresponding to 10 mg/mL protein immobilised on the surface). For ArgD, a 1 mM PLP injection after immobilization increased the absolute baseline immobilisation response by ≈1100 RU. This was done to compensate for loss of PLP in the low pH coupling buffer during immobilisation. The baseline throughout the screening experiment remained at the post-PLP injection level (≈8570 RU), indicating that PLP was not lost during the experiment.

The immobilised ArgC protein was tested first using dilution series of NADPH and NADP^+^ (19 μM to 2.5 mM), and a clear dose response suggested that predominantly, the enzyme had not been immobilised in an orientation that occluded the active site. Similarly, ArgD was tested using a dilution series of L-glutamate, N-acetylornithine and L-ornithine (19 μM to 2.5 mM).

The fragment hits obtained previously were injected at a concentration of 1 mM. The sensorgrams for all the analytes were inspected visually to exclude fragments with either no discernible response or a “sticky” profile from further analysis. The binding level was calculated using the T200 software and adjusted for molecular weights of the analytes. For ArgC, 22 fragments with a binding level response ≥20 RU were shortlisted, whereas in the case of ArgD, 20 fragments with a response ≥10 RU were shortlisted for crystallographic validation. Fragments were thereafter described either as SPR ‘positive’ or ‘negative’.

### 3.3. Crystallographic, biophysical and biochemical validation

The hits obtained for ArgB, ArgC, ArgD and ArgF were then soaked in crystals of the respective protein and X-ray diffraction data was collected. Data collection and refinement statistics for all structures are available in Table S3.

#### 3.3.1. ArgC

Crystal structures were obtained for the ArgC apoenzyme (PDB: 7NNI) and the NADP^+^-bound holoenzyme (PDB: 7NNQ) (Figure 2A). Binding of NAPD causes significant structural changes in two loops of the protein that move from a closed to open conformation (Figure 2A). Additionally, structures were solved with 4 fragment binders, occupying either of the two distinct pockets: the substrate-binding (Figure 2C and D) and the NADP(H)-binding pockets (Figure 2E and F). Fragments NMR322 (5-Methoxy-3-indoleacetic acid) and NMR 571 (Xanthene-9-carboxylic acid) were observed in the substrate-binding pocket (PDB: 7NOT and 7NNR respectively) (Figure 2C and D). Both NMR322 and NMR571 engaged side chains of residues His217 and Tyr211 through hydrogen bonds. Both residues are predicted to stabilise the acyl-enzyme intermediate during catalysis. NMR322 also made an H-bond interaction with ser186 and gly187. As compared to NMR571, NMR322 binds deeper in the pocket (Figure 2C and D).

**Figure 2:**
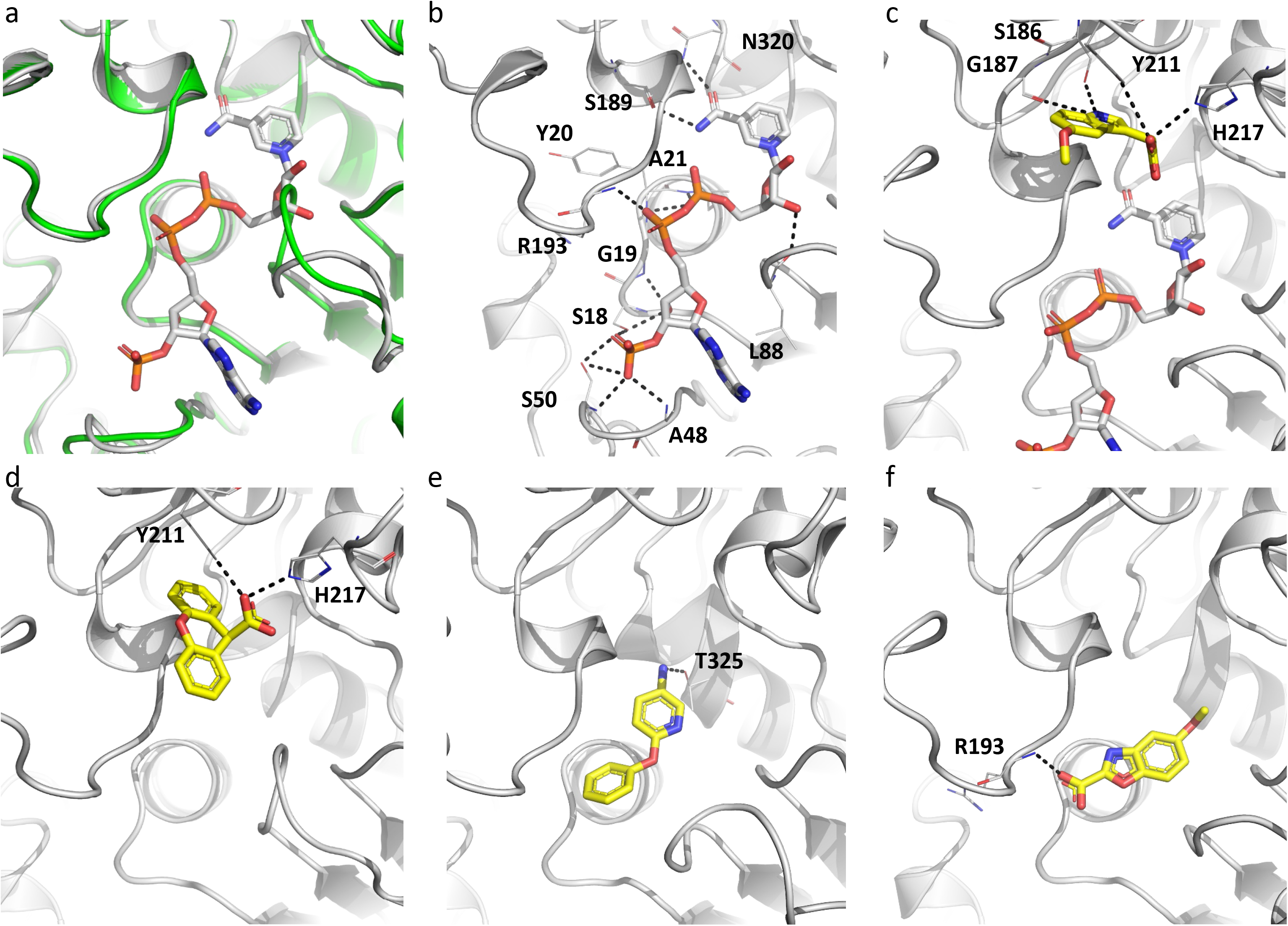
(A) X-ray crystal structure of ArgC apoenzyme superposed with the NAPD bound holoenzyme. X-ray crystal structures of ArgC in complex with fragments NMR322 (a) and NMR571 (b) binding to the substrate site and NMR401 (c) and NMR863 (d) binding to the co-factor site. Hydrogen bonds are represented by black dashed lines.

Fragment NMR401 (6-phenoxy-3-pyridinamine) and NMR863 (5-methoxy-1,3-benzoxazole-2-carboxylic acid) were observed in the ribosylnicotinamide and pyrophosphate regions of the NADP(H)-binding pockets (PDB: 7NPJ and 7NPH respectively) and the majority of the interactions between the protein and these two fragments are hydrophobic or π-interactions. Both fragments form only a single hydrogen bond interaction with the thr325 side chain in case of NMR401 (Figure 2E) and with the arg193 backbone amine in the case of NMR863.

Enzymatic assays revealed that 2 mM NMR322 inhibited ArgC activity by 45% whereas 2 mM NMR571 caused a 37% inhibition. 2 mM NMR401 inhibited ArgC activity by 10% whereas 2 mM NMR863 caused a 12% inhibition. Although the thermal shifts obtained for NADP(H)-binding pocket fragments NMR401 and NMR863 were higher than those for substrate-binding pocket fragments NMR322 and NMR571, the SPR binding response for the latter was better and positively correlated with percentage inhibition of enzymatic activity (**Table 2**).

#### 3.3.2. ArgD

The first crystal structure of the ArgD holoenzyme from *M. tuberculosis* obtained (Figure 3A) showed that the protomer has three domains: the smaller N-terminal segment (residues 7 to 74), the relatively larger C-terminal domain (residues 286 to 396), and the central PLP-binding domain (residues 85 to 273), which is also the largest and has a Rossmann-like overall fold (Figure 3B). The prosthetic group PLP is covalently linked to Lys253 via an aldimine linkage. ArgD is a dimeric enzyme like other members of the class III δ-aminotransferase family (38); the active sites are interfacial, and residues of both protomers contribute to the active site architecture (Figure 3A) (PDB: 7NN1).

**Figure 3:**
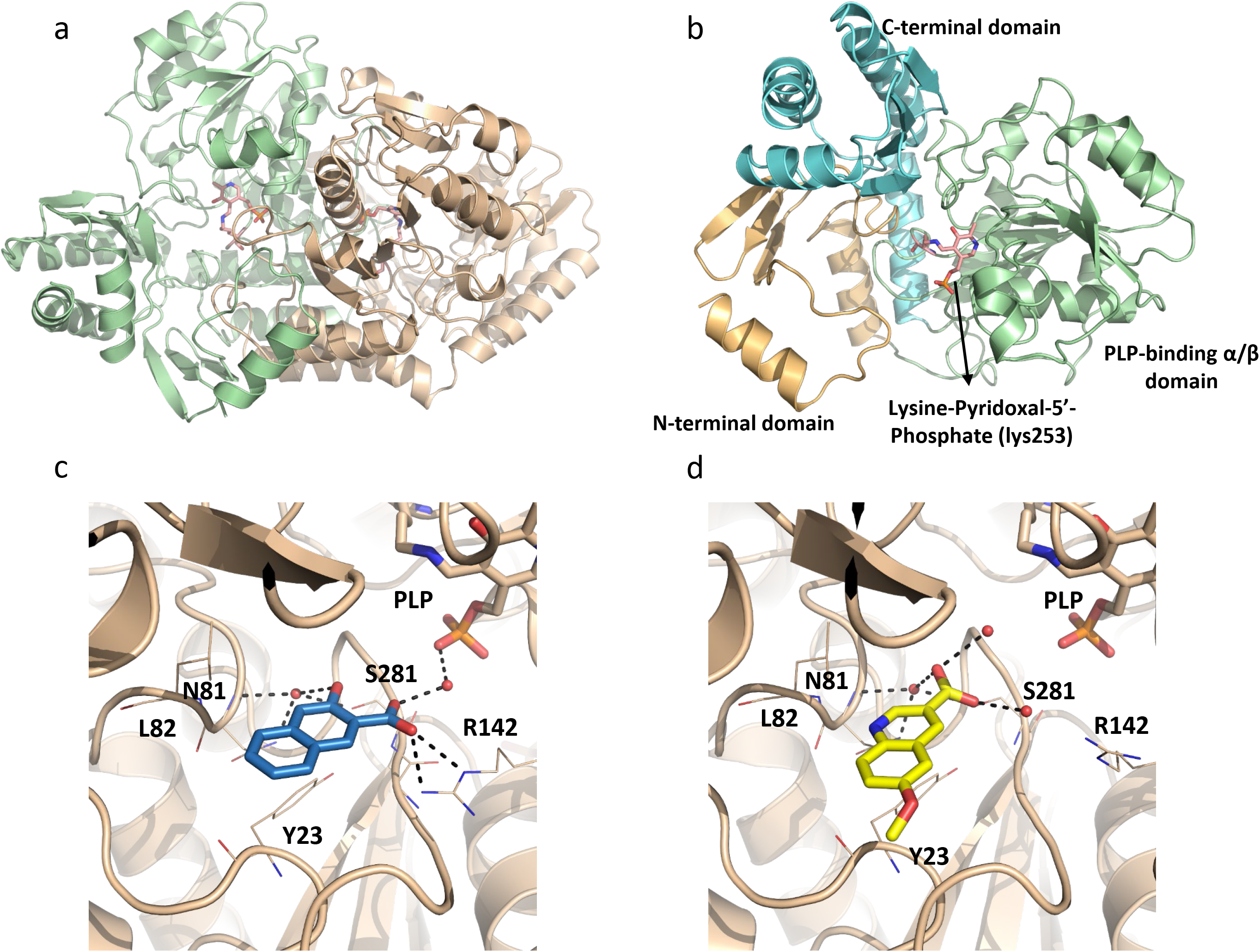
(a) X-ray crystal structure of *M. tuberculosis* ArgD showing the dimer. Each protomer of the dimer is highlighted in a different colour. The ArgD protomer comprises of three domains shown in gold (N-terminal domain), green (PLP-binding α/β domain) and cyan (C-terminal domain) (b). X-ray crystal structures of ArgD in complex with fragments NMR608 (c) and NMR868 (d). Hydrogen bonds are represented by black dashed lines.

Fragment NMR608 (3-Hydroxy-2-naphthoic acid) was observed occupying all four binding sites (4 chains in the ASU). It binds closer to the distal phosphate group of PLP than the proximal ring system and the internal aldimine bond, where catalysis actually occurs. However, it makes, through its acid group, hydrogen bond interactions with Arg142, which could be the key residue involved in substrate binding based on homology with AstC, and with an highly coordinated water through the hydroxyl group (Figure 3C). While the pocket itself has depth, the fragment is more solvent-exposed than the PLP molecule. While NMR608 was the only hit validated by all techniques the fragment library contained a compound with similar structure (NMR868) that had only a melting shift of 0.9 C and therefore wasn’t selected initially for further testing. However, soaking of this fragment revealed that it binds in almost the same position as NMR608 (Figure 3D). Nevertheless, The orientation of the acid group is slightly different in NMR868 and it no longer interacts directly with arg142. In fact this fragment only has hydrogen bond interactions with solvent molecules and it is only observed in one molecule in the ASU out of four, most likely reflecting a lower affinity (PDB: 7NNC).

Aminotransferases are often assayed for the reverse reactions they catalyse because in most cases substrates for the forward reaction are not commercially sold. The ArgD holoenzyme can also use N-acetylornithine (NAO) and α-ketoglutarate (α-KG) to generate N-acetyl-γ-glutamyl-semialdehyde and glutamate. The semialdehyde product spontaneously cyclises into Δ^1^-pyrroline-5-carboxylic acid and can react with the reagent 2-aminobenzaldehyde to yield a dihydroquinazolinium compound (bright yellow colouration) that absorbs at 440 nm (39, 40). This assay was employed to assess the effect of the fragment hits on on ArgD. NMR608 exhibited very weak activity with only 31% and 16% inhibition observed at 4 and 2mM respectively. This is consistent with the crystal structures where the fragment is highly exposed to the solvent.

#### 3.3.3. ArgF

Crystal structures of apo ArgF and of ArgF in complex with the natural ligand carbamoyl phosphate were initially obtained (PDB: 7NNF and 7NNV respectively). Carbamoyl phosphate interacts with ArgF through hydrogen bonds with the side chains of ser50, thr51, arg52, arg101, his128 and gln131 but also with the backbone amines of thr51, arg52 and thr53 and the carbonyl of cys264 (Figure 4A). The residues ser50, thr51, arg52 are at the N-terminus of α-helix 2 and the phosphate group of carbamoyl phosphate sit at the positive pole of the helix. Binding of carbamoyl phosphate to ArgF slightly displaces α-helix 2 when compared to the apo structure (Figure S2A). Soaking ArgF with fragment hits yielded eight crystals structures. All fragments occupied a site at the interface between two protomers of the ArgF trimer but not all sites are equally occupied by all fragments (Figure 4B-E and S2B-F). This site contacts directly with the α-helix 2 and sits between this helix and α-helix 3 of the opposing protomer. The site is formed by residues thr51, arg52, phe55 of α-helix 2, leu265, ala289, arg292 of protomer 1 and ile45, ser75, thr76, leu78, glu82, thr87, leu91 and tyr94 of protomer 2. These residues form a cavity that opens to the carbamoyl phosphate binding site. Binding of fragments at this site, slightly shifts the position of α-helix 2 by 1.9 Å, when compared to the carbamoyl phosphate structure, and also affects the conformation of arg52 which is involved in carbamoyl phosphate binding. Fragments were found to bind in two distinct sub sites across the main binding site and could be divided in three different groups based on their mode of binding. NMR007, 078, 464 and 502 occupied the top area of the site (subsite 1) and contacted with a loop of protomer 2 that is composed by residues asp72 to leu84 that covers the site (Figure 4C and S2B-D). NMR801, the single representative of this group sits at the bottom of the site (subsite 2) between α-helix 2 of protomer 1 and α-helix 3 of the opposing protomer (Figure 4D). NMR288, 812 and 817 have two molecules binding at this site, with one molecule occupying each subsite (Figure 4E and S2E-F). All fragments keep α-helix 2 in a position close to the apo structure or move further away from the position this helix occupies when carbamoyl phosphate is bound, albeit very slightly. A *K*_d_ value could only be determined for NMR007 and NMR812 and both fragments showed affinities worse than 100 μM (Table 4 and Figure S3).

**Figure 4:**
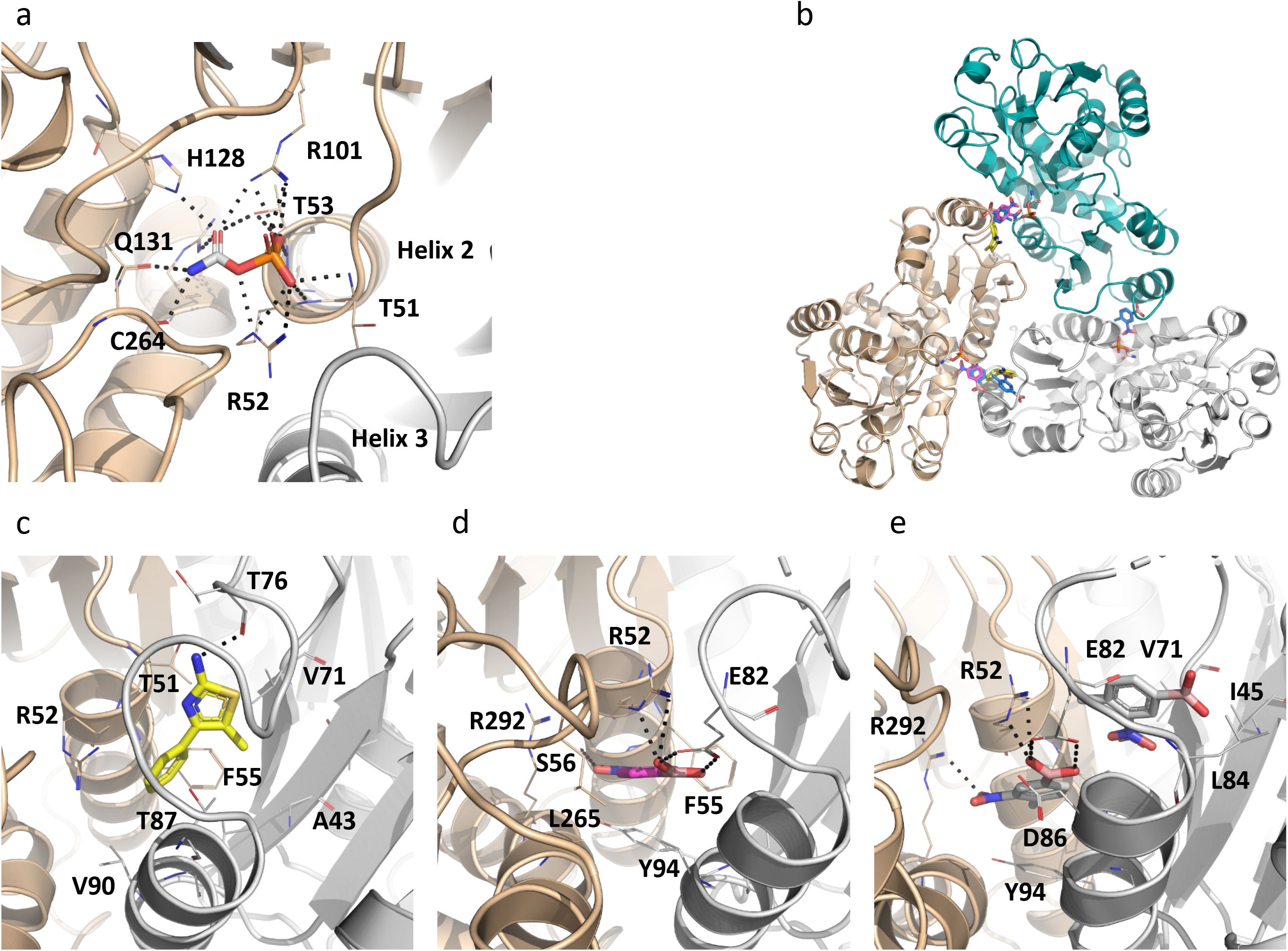
(A) X-ray crystal structure of *M. tuberculosis* ArgF in complex with carbamoyl phosphate. Hydrogen bonds are depicted as black dashed lines. Two of the protomers of the trimer are visible and are coloured differently. (B) Structure of the ArgF trimer bound with fragments at interfacial site. Three X-ray crystal structure of ArgF in complex with different fragments were superposed with the apo structure to create this figure. (C) X-ray crystal structures of ArgF in complex with NMR007 (C), representing the group that binds to subsite 1, NMR801 (D), the single representative of the group that binds to subsite 2 and NMR812 (E), representing the group of fragments that bind to both subsites, with hydrogen bonds depicted as black dashed lines.

#### 3.3.4. ArgB

We obtained crystal structures for ArgB in the apo form and with the natural ligands N-acetyl-glutamate (NAG) and L-arginine (PDB: 7NLF, 7NLN, and 7NLO respectively) (Figure 5A). The thirteen crystal structures of ArgB with fragments in the absence of natural ligands show that all the compounds were unexpectedly bound to a hydrophobic cavity at the interface between two protomers with three of these sites present in the ArgB hexamer (Fig. 5B-D and Figure S4). L-canavanine, the guanidinooxy structural analogue of L-arginine, bound to the same site of ArgB as L-arginine and induced conformational changes similar to those induced by L-arginine (Figure S5). The interfacial site is composed by ala124, val125, gly126, ile127, asp131, ala132, leu134, ala164, met165, leu168 and arg173 and is mostly hydrophobic in nature (Figure S6). Due to the nature of this new site, the interactions between the compounds and the protein are essentially hydrophobic, with residues leu168 and val125 interacting with NMR711 (PDB: 7NNB) while for NMR446 (PDB: 7NLX) ile127 is also involved in the hydrophobic interactions (Fig. 5C and D). Carbon-π interactions are also formed with leu168 for both compounds (Fig. 5C and D). Finally, weak hydrogen bonds are also present between asp167, leu168 and val125 for NMR711 while NMR446 interacts with ile127 and leu168 via hydrophobic contacts (Fig. 5C and D). Furthermore, this site is symmetrical and sits at a 2-fold crystallographic symmetry axis with each compound clearly presenting two binding conformations (Figure 5E and S4). The two compounds (NMR446 and NMR711) also share structural and binding features with a trifluoromethyl group occupying the same position at the binding site.

**Figure 5:**
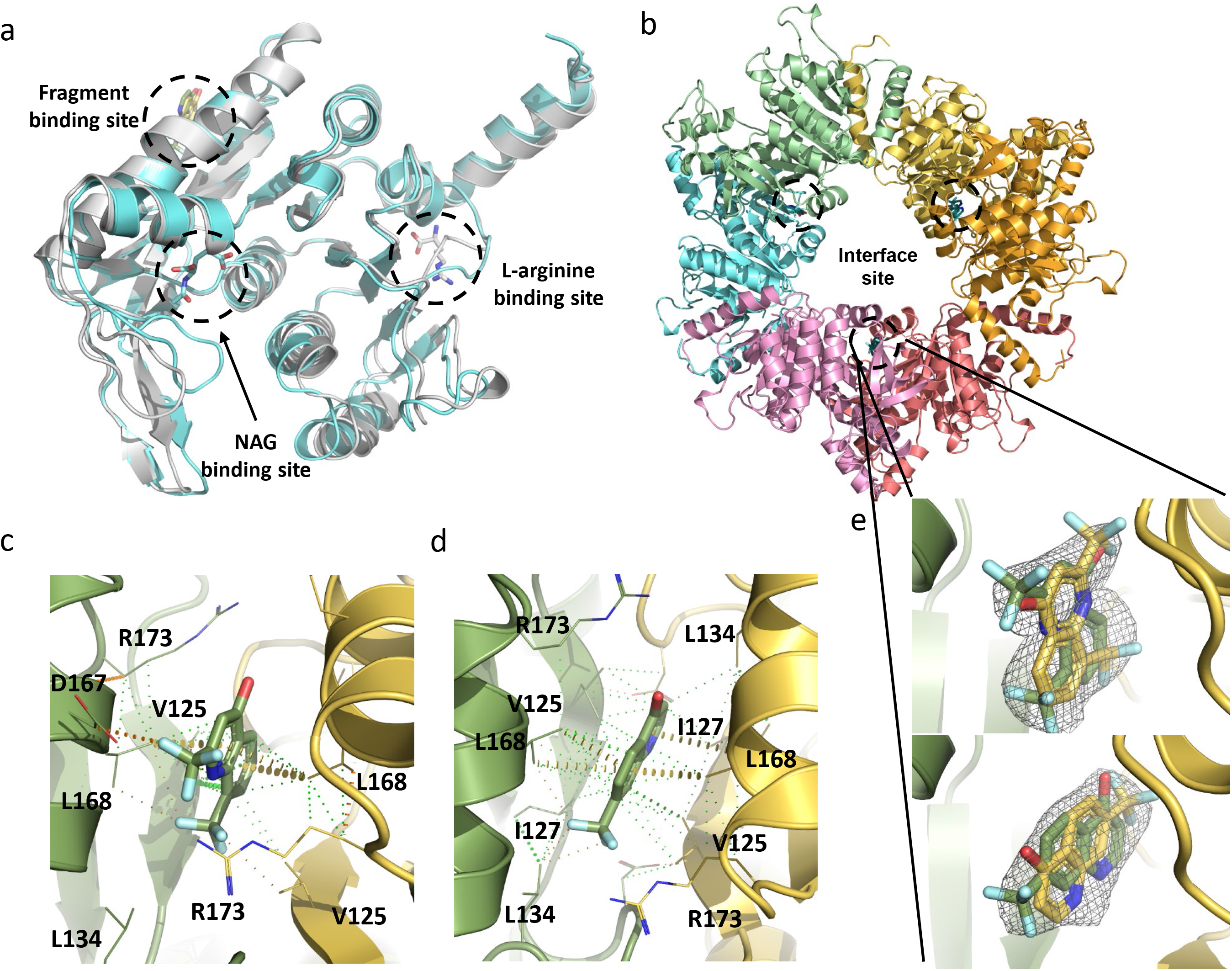
(A) Overlap of X-ray crystal structures of protomers of Apo-ArgB, ArgB co-crystallized with N-acetyl glutamate (NAG), and ArgB co-crystallized with L-arginine. (B) Structure of the ArgB hexamer with fragments bound at the interfacial site of two protomers. Each ArgB protomer is coloured differently. X-ray crystal structures of ArgB in complex with NMR711 (c) and NMR446 (d). Hydrophobic interactions are depicted in green dots, weak hydrogen bonds in orange dots and carbon-π interactions in yellow disks. Only one binding conformation is shown for clarity in both panels. [Fo - Fc] “Omit maps” of NMR711 (e) NMR446 (f) contoured at 1.5σ. These maps were generated with using the phases from the final model. The two adopted conformations are shown for both compounds.

Enzymatic assays show that, of the 15 compounds, NMR711 [2,8-bis(trifluoromethyl)-1H-quinolin-4-one] and NMR446 [8-(trifluoromethyl)-1H-quinolin-4-one] were the best ArgB inhibitors, with an IC_50_ of 366 and 707 μM, respectively (Fig. 6A and Table 1). The natural allosteric regulator L-arginine and its analogue L-canavanine (Fig. 6A) have IC_50_ values of 186 μM and 1.46 mM respectively. Additionally, ITC experiments showed that compounds NMR711 and NMR446 bind to ArgB with a KD of 7.7 and 23 μM, respectively (Table 1 and Figure S7), whereas L-arginine and L-canavanine showed complex binding curves that can only be fitted to a sequential binding model showing a cooperative interaction with the different protomers of the hexamer (Table 1 and Figure S7). NMR competition assays revealed that compounds NMR711 and NMR446 are not competitive with any of the natural ligands (ATP, NAG, and L-arginine), with enzymatic assays also demonstrating the non-competitive nature of the inhibition of both fragments. This confirms that the results from X-ray crystal structures are not an artefact and that the fragment-binding site is indeed a new allosteric site (Fig. 6B and S8). The observation that there are no conformational changes in the crystal structures of ArgB with either fragment at the obtained resolution (2-2.5 Å) may be due to constraints arising from crystal packing. Bioinformatics analysis showed that this site is conserved in mycobacterial species and also in closely related actinobacteria, such as Nocardia (Figure S6B). Nevertheless, it is clear that binding of these compounds, similar to L-arginine binding, causes changes in the energy landscape of the protein that result is allosteric inhibition of the catalytic reaction.

**Figure 6:**
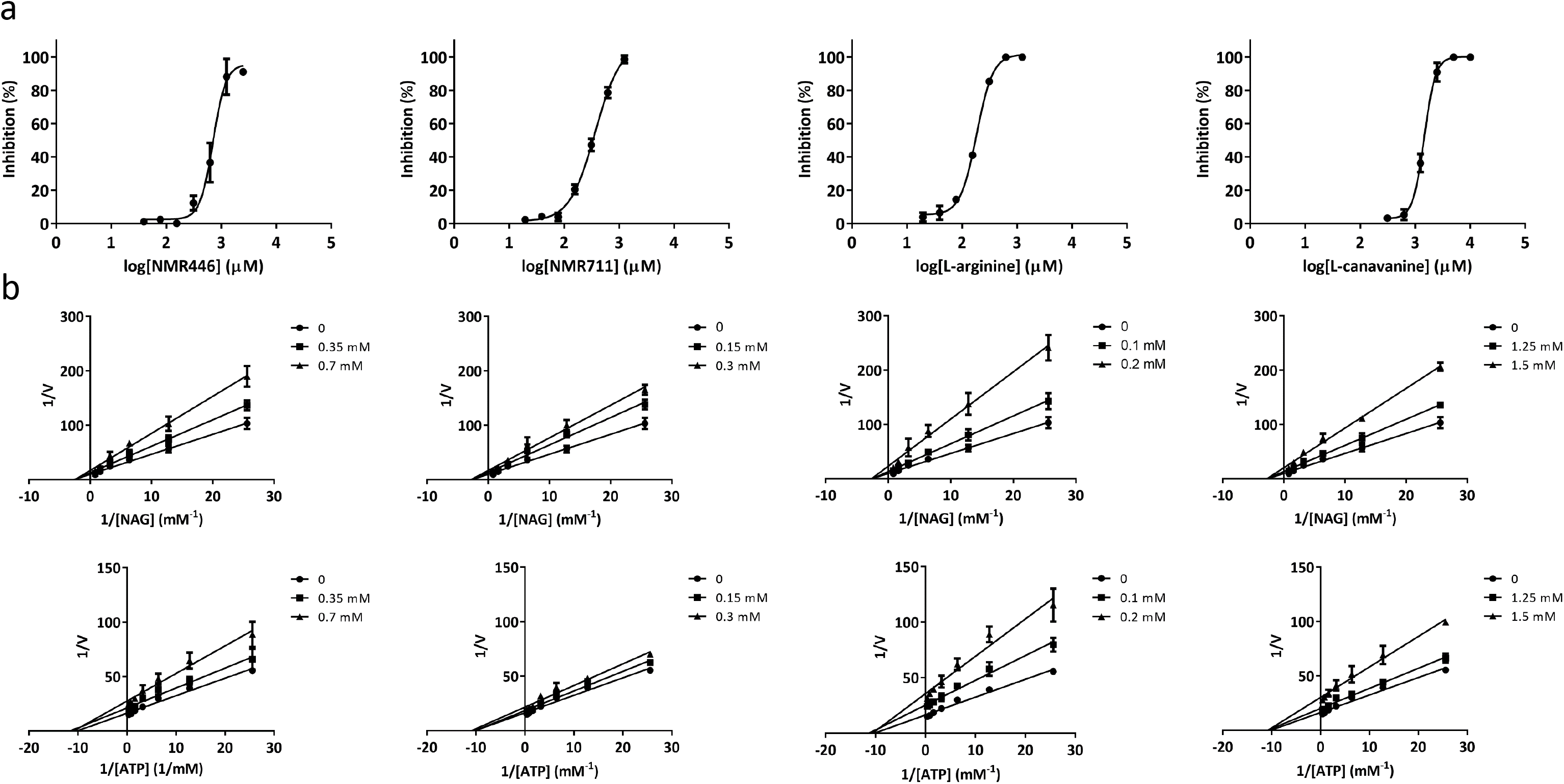
ArgB inhibition by arginine analogs and allosteric fragment inhibitors. (a) Inhibition of ArgB activity by NMR711, NMR446, L-arginine and L-canavanine. (b) Lineweaver-burk plots for NMR711, NMR446, L-arginine and L-canavanine. Average of replicates and standard deviation are ploted (n=3).

### 3.4. Effect of ArgB inhibitors in *M. tuberculosis* growth

Considering the four enzymes screened of the arginine biosynthesis pathway, ArgB hits exhibited higher potency by far, with NMR711 and NMR446 being selected to assess their effect on *M. tuberculosis* together with L-canavanine.

The ability of these compounds to inhibit *M. tuberculosis* growth was examined by measuring their minimum inhibitory concentrations (MICs) in the absence or presence of arginine (1 mM). All the compounds inhibited the growth of *M. tuberculosis* H37Rv and Δ*argB*-c in media without arginine compared to no drug control (Figure 7). MICs for NMR711, NMR446 and L-canavanine were 25-50, >200 and 50 μg/ml against H37Rv and Δ*argB*-c (Figure 7). However, when arginine (1 mM) was present in the media, compound NMR446 and L-canavanine had no inhibitory activity against H37Rv, ΔargB-c and, Δ*argB*. This indicates that these compounds are indeed specifically inhibiting *M. tuberculosis* arginine biosynthesis (Figure 7). In contrast, the more promiscuous NMR711 was inhibitory for all the above strains in the presence or absence of arginine, suggesting that NMR711 may target additional proteins (Figure7).

**Figure 7:**
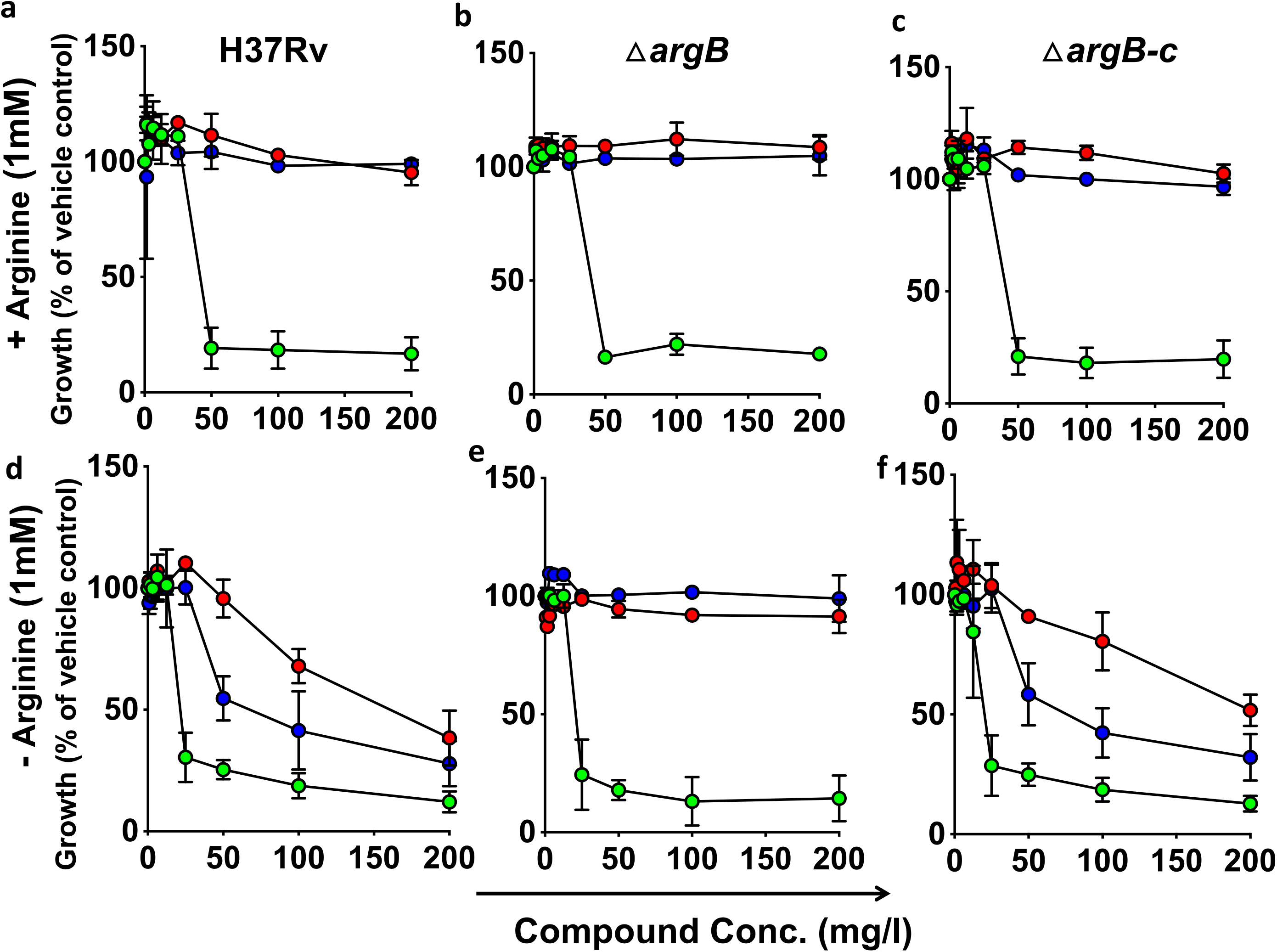
Dose response curves of inhibitor compounds for inhibition of *M. tuberculosis* growth (measured as optical density at 600nm) in the presence (a-c) or in the absence (d-f) of 1mM L-arginine. Data is represented as percentage growth of *M. tuberculosis* strains in the presence of different concentrations of the inhibitor compared to growth in the presence of just vehicle control (no drug). H37Rv (a,d), complemented Δ*argB* (Δ*argB*-c; b,e), and Δ*argB* (c,f). Data are representative of one of three independent experiments. Error bars, mean s.d. (n = 3). Compound 1 (green circles), Compound 2 (red circles) and L-Canavanine (blue circle).

Using the FBDD approach, we have discovered inhibitors that bind to a new allosteric site in ArgB, which has very different properties than that of the active site and L-arginine binding sites, thus opening new possibilities for drug discovery by targeting ArgB. For fragment-sized molecules, both compounds reported in this work bind tightly and allosterically to ArgB and have growth inhibitory activity against *M. tuberculosis,* suggesting that they have the potential to provide a framework for developing larger and higher affinity molecules against the ArgB protein.

## 4. Discussion

The arginine biosynthesis pathway has been established as a good target for anti-TB drug discovery (10, 14). Arginine deprivation in *M. tuberculosis* induced by knocking out *argB* and *argF* results in both *in vitro* and *in vivo* sterilisation of *M. tuberculosis*, without the emergence of suppressor mutants (10). However, from a pathway with eight enzymes, only ArgJ has been explored in a drug discovery campaign and all other enzymes of the pathway, prior to this work, were yet to be assessed in their potential as suitable targets for drug discovery.

Fragments are potent chemical tools that can efficaciously explore the surface of proteins for new binding sites and their chemical space, even with small libraries of a few hundreds of compounds and can therefore be employed to assess the ligandability of protein targets (18, 41). Therefore, this approach was employed to assess the ligandability of ArgB, ArgC, ArgD and ArgF, to identify potential starting points for fragment development.

We have screened a fragment library of 960 small compounds (MW 150-300 Da) initially using DSF and employed ligand-based NMR, SPR, ITC, biochemical assays and X-ray crystallography to validate the hits. Due to the nature of FBDD a hit is only considered validated when an X-ray crystal structure is obtained. For all the proteins in this work, hits were found and eventually validated by X-ray crystallography. ArgB had the highest number of X-ray validated hits with a total of fourteen, followed by ArgF with eight, ArgC with four and ArgD with two. Interestingly, in the case of ArgB and ArgF, all the fragment hits were binding to an interfacial site, which in the case of ArgB was confirmed to have functional implications. In the case of ArgF, its close proximity to the active site shows potential to develop compounds that can anchor at the interfacial site and then extend towards the active site, thus inhibiting the enzyme. Similarly for ArgJ, the only enzyme of the pathway with known inhibitors prior to this work, the inhibitors were also found to bind to an interfacial allosteric site (14). Our results further show that in the case of ArgC there are two possible strategies to develop inhibitors, with one targeting the cofactor binding site and the other the substrate binding site. It is not clear at this point which strategy has the highest potential to result in potent inhibitors. Another consideration to take into account is the level of homology of these enzymes with the human orthologue. While ArgB and ArgC do not have a human orthologue, ArgD and ArgF do and the *M. tuberculosis* enzymes have identities of 36 and 41% with the human orthologues, respectively. Nevertheless, while the ArgF active site is conserved, the interfacial site of ArgF contains several differences that raise the prospect of developing specific inhibitors for the *M. tuberculosis* enzyme. For ArgD selectivity might be more difficult to achieve since many of the active site residues are conserved in comparison with the human cytoplasmic and mitochondrial enzymes.

Due to the potency of the best fragments against ArgB, we tested them for their ability to inhibit *M. tuberculosis* growth together with L-canavanine. Remarkably, NMR446 and L-canavanine not only inhibited *M. tuberculosis* growth, but were also found to act on-target despite the potential promiscuity of such small compounds, with both becoming inactive after the addition of L-arginine to the media.

Despite these promising results, the interfacial site of ArgB might be the hardest of all sites found in this study to develop small molecule inhibitors. The intrinsic highly hydrophobic nature of the site together with very few opportunities to engage in hydrogen bonds and other polar contacts creates difficulties in rationalizing what modifications could improve the potency of the compounds. Furthermore, the fact that we cannot observe conformational changes in any of the structures with fragments bound to ArgB may be due to constraints arising from crystal packing and thus these structures may not completely represent what happens in solution. It is however also possible that binding to this site does not cause visible conformational changes but still alters the energy landscape of the intramolecular pathways involved in the catalytic cycle. We cannot currently determine which of these two hypotheses is correct.

In conclusion, using a fragment-based approach, we have discovered inhibitors that bind to novel sites in ArgB and ArgF and to the active sites of ArgC and ArgD, which in case of ArgB show on target activity against *M. tuberculosis*. The data presented here clearly shows that there is scope to target at least ArgC and ArgF with dedicated drug discovery programs and we propose these two as the best candidates for future drug discovery work.

## Abbreviations

TB: tuberculosis
FBDD: Fragment-based drug discovery
DSF: Differential scanning fluorimetry
ASU: asymmetric unit
SPR: Surface plasmon resonance
NMR: Nuclear magnetic resonance
ITC: Isothermal titration calorimetry

## Acknowledgements

This work was funded by Bill and Melinda Gates Foundation HIT-TB (OPP1024021) and SHORTEN-TB (OPP1158806). PA was funded by a Gates Cambridge Scholarship. TLB is funded by the Wellcome Trust (Wellcome Trust Investigator Award 200814_Z_16_Z: RG83114). The authors would like to thank the Diamond Light Source for beam-time (proposals mx14043 and mx18548).

