## Supplements for "A Fragment-based approach to assess the ligandability of ArgB, ArgC, ArgD and ArgF in the L-arginine biosynthetic pathway of *Mycobacterium tuberculosis*"

* To whom correspondence should be addressed

### Methods

**Table S1:** Buffers used for protein purification and storage

<table>
<tr><th>Protein</th><th>Buffer A</th><th>Buffer B</th><th>Buffer C</th><th>Storage buffer</th></tr>
<tr><td>ArgB</td><td>25 mM HEPES pH 7.5<br>500 mM NaCl<br>20 mM imidazole</td><td>25 mM HEPES pH 7.5<br>500 mM NaCl<br>500 mM imidazole</td><td colspan="2">25 mM HEPES pH 7.5<br>200 mM NaCl</td></tr>
<tr><td>ArgC</td><td rowspan="2">20 mM Tris-HCl pH 7.4<br>500 mM NaCl<br>20 mM imidazole</td><td rowspan="2">20 mM Tris-HCl pH 7.4<br>500 mM NaCl<br>500 mM imidazole</td><td>20 mM Tris-HCl pH 7.4<br>500 mM NaCl</td><td>5 mM Tris-HCl pH 7.4<br>50 mM NaCl</td></tr>
<tr><td>ArgD</td><td colspan="2">50 mM Tris-HCl pH 7<br>100 mM NaCl</td></tr>
<tr><td>ArgF</td><td>50 mM Sodium Phosphate pH 7.4<br>500 mM NaCl<br>20 mM imidazole</td><td>50 mM Sodium Phosphate pH 7.4<br>500 mM NaCl<br>500 mM imidazole</td><td colspan="2">50 mM Sodium Phosphate pH 7.4<br>100 mM NaCl</td></tr>
</table>

### Results

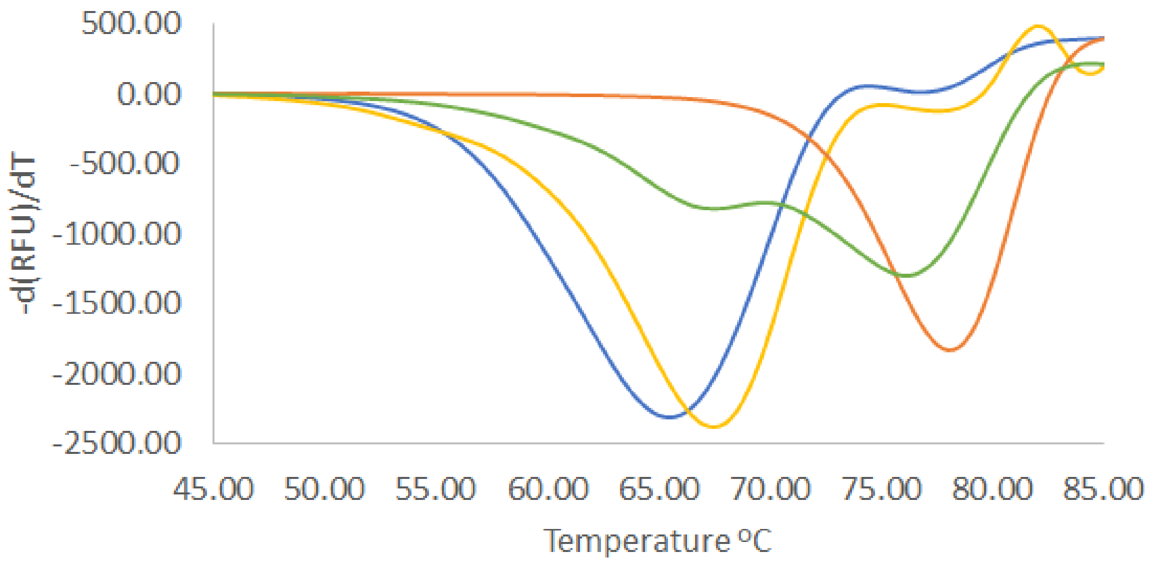


**Figure S1**: DSF profiles of ArgD showing PLP-unsaturated ArgD (blue), in the presence of 1 mM PLP (orange), and in the presence of two different fragments (yellow and green). PLP-unsaturated ARgD presents two melting peaks, the largest one representing the protein populations without PLP-bound in the dimer or with only protomer containing PLP. The smaller peak has similar melting temperature to the PLP saturated enzyme and likely represents a population where both protomers contain PLP. While the fragment with the green profile alters the melting profile of the protein with the major peak now having a melting temperature

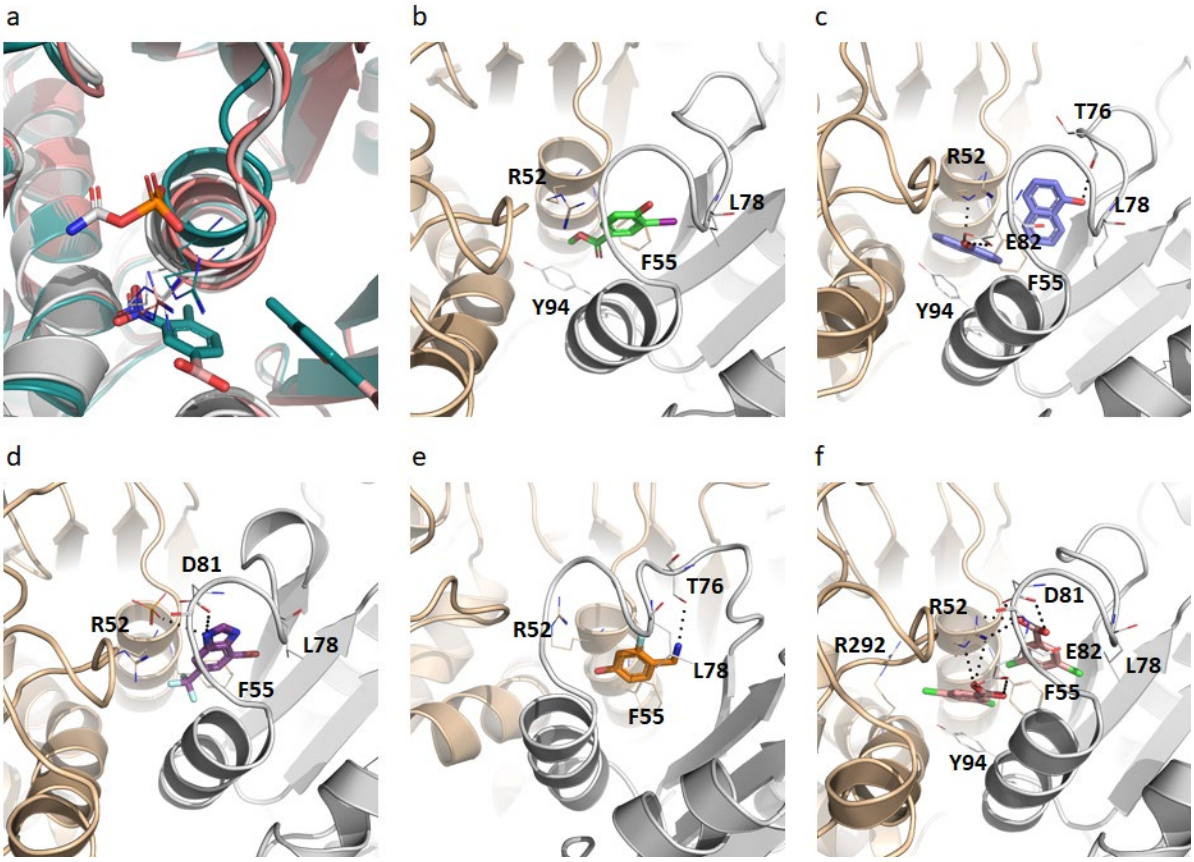


**Figure S2**: (a) Superposition of ArgF apo structure in pink with the carbamoyl phosphate bound structure in white and NMR817 structure in teal, showing movement of helix 2 upon binding of ligands. The distance the α carbon of arg52 is 1.3 Å between the apo and carbamoyl phosphate structures and 1.9 Å between the NMR817 and carbamoyl phosphate structures. X-ray crystal structures of ArgF in complex with compounds NMR078 (b), NMR288 (c), NMR464 (d), NMR502 (e), NMR812 (f). Black dashed lines denote hydrogen bonds. Each protomer of the interfacial site is coloured differently.

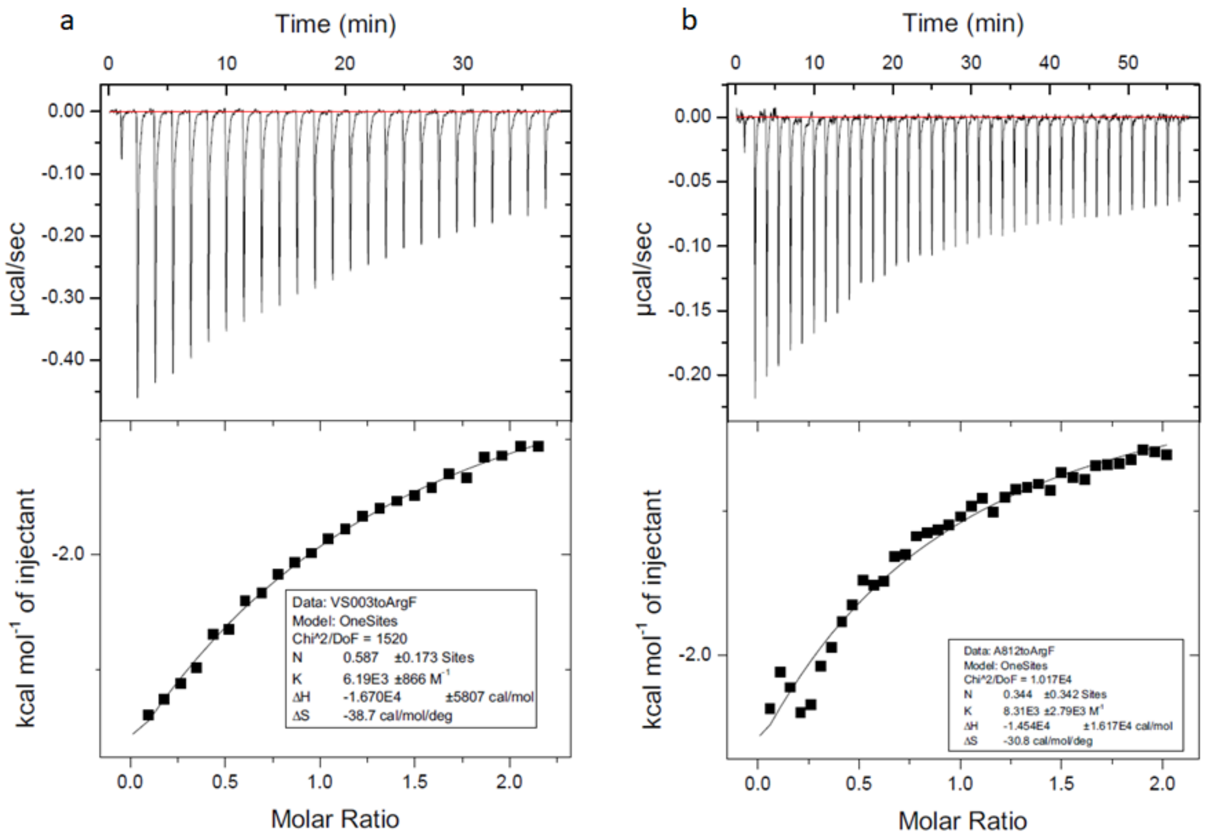


**Figure S3**: ArgF ITC titrations for NMR007 (a) and NMR812 (b).

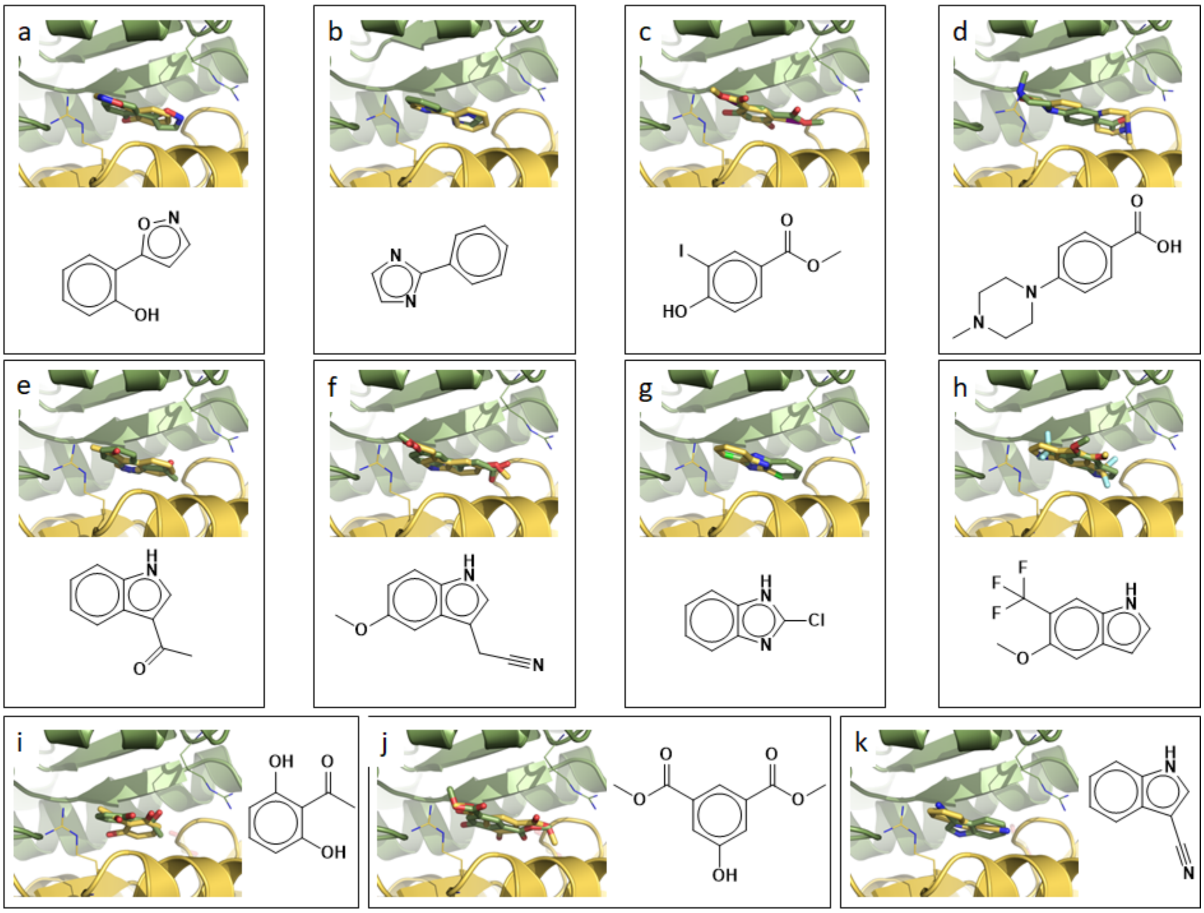


**Figure S4:** X-ray crystal structures of ArgB in complex with compounds NMR026 (A), NMR043 (B), NMR078 (C), NMR082 (D), NMR314 (E), NMR323 (F), NMR462 (G), NMR469 (H), NMR582 (I), NMR612 (J) and NMR617 (K). R173 is shown in every panel delimitating the binding site on both protomers. As the site is symmetrical and sits at a 2-fold crystallographic symmetry axis each fragment exhibits two binding conformations. Chemical structures of the compounds are shown within each panel for clarity.

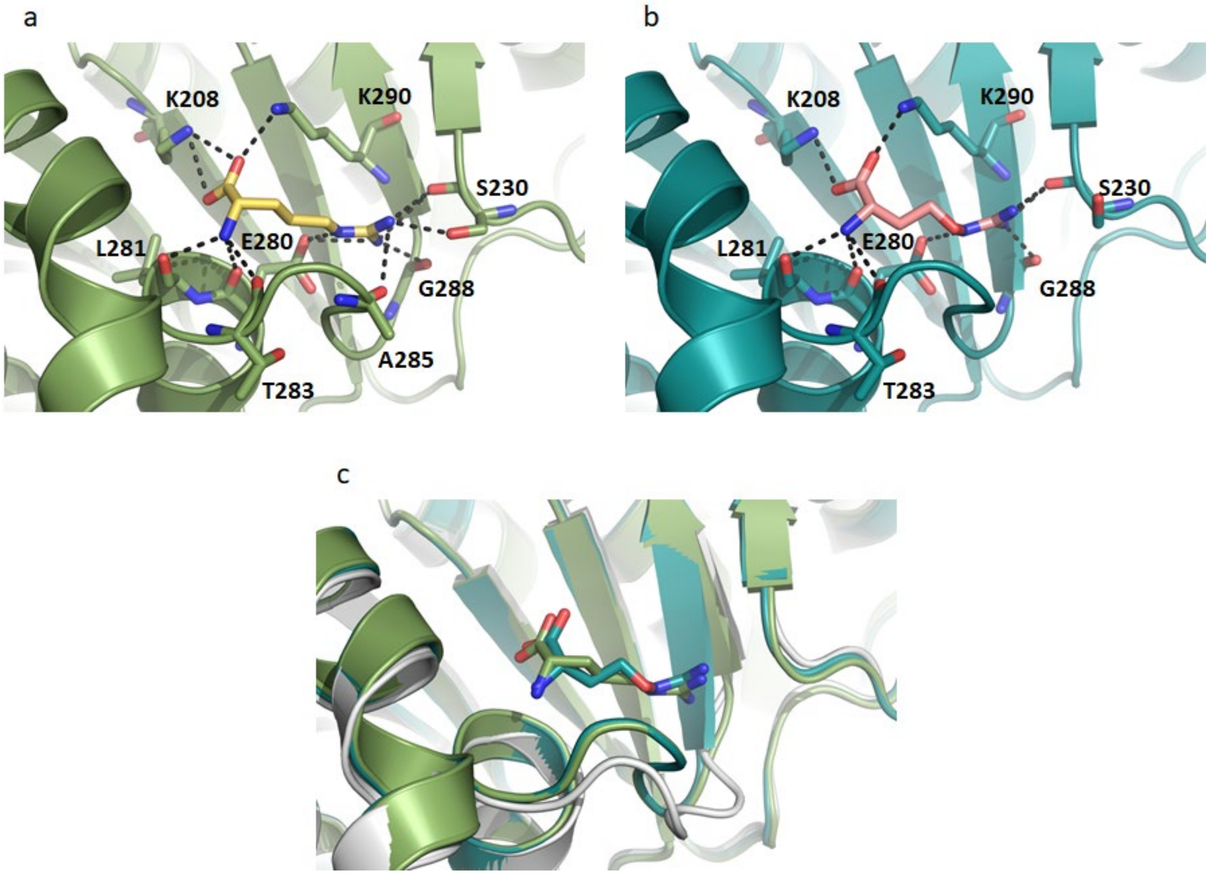


**Figure S5:** X-ray crystal structures of ArgB in complex with L-arginine (a) and L-canavanine (b). Hydrogen bonds between ligands and residues are shown. (c) Superposition of ArgB Apo structure (white) with ArgB in complex with L-arginine (green) and L-canavanine (blue) illustrating the conformational changes induced by ligand binding. L-arginine and L-canavanine binding to ArgB induce similar conformational changes.

a

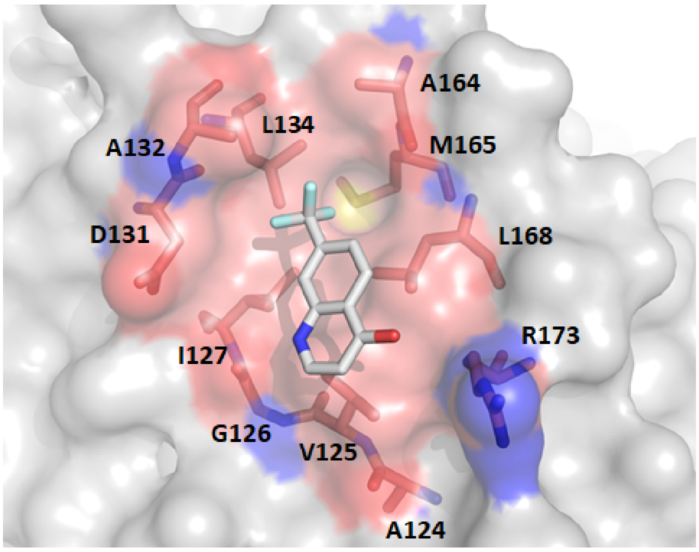


b

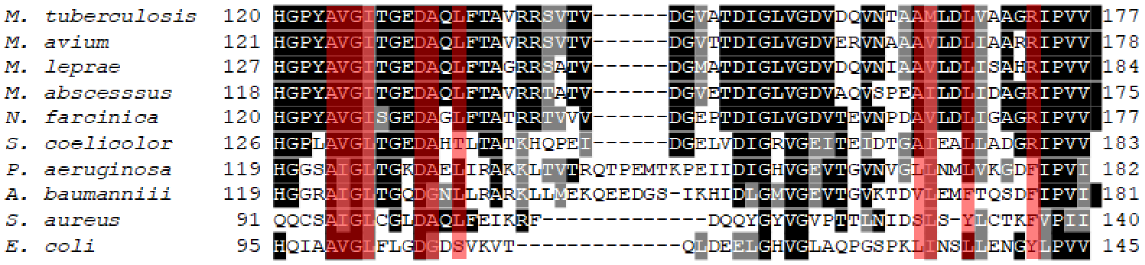


**Figure S6**: (**A**) Crystal structure of ArgB in complex with NMR446 with interface site residues highlighted in red. Only one conformation of the ligand and one protomer are shown for clarity. (**B**) Alignment of ArgB sequences from *Mycobacterium tuberculosis*, *Mycobacterium avium*, *Mycobacterium leprae*, *Mycobacterium abscessus*, *Nocardia farcinica*, *Streptomyces coelicolor*, *Pseudomonas aeruginosa*, *Acinetobacter baumannii*, *Staphylococcus aureus* and *Escherichia coli* was performed using clustal omega (1). The alignment shows high conservation of residues at the interface site in mycobacteria and closely related species. Non-actinobacterial species (*P. aeruginosa*, *A. baumannii*, *S. aureus* and *E. coli*) show lower conservation. Residues forming this site are highlighted in red. ArgB of *E. coli* and *S. aureus* are dimeric as they lack the N-terminal helix that allows the formation of the hexamer.

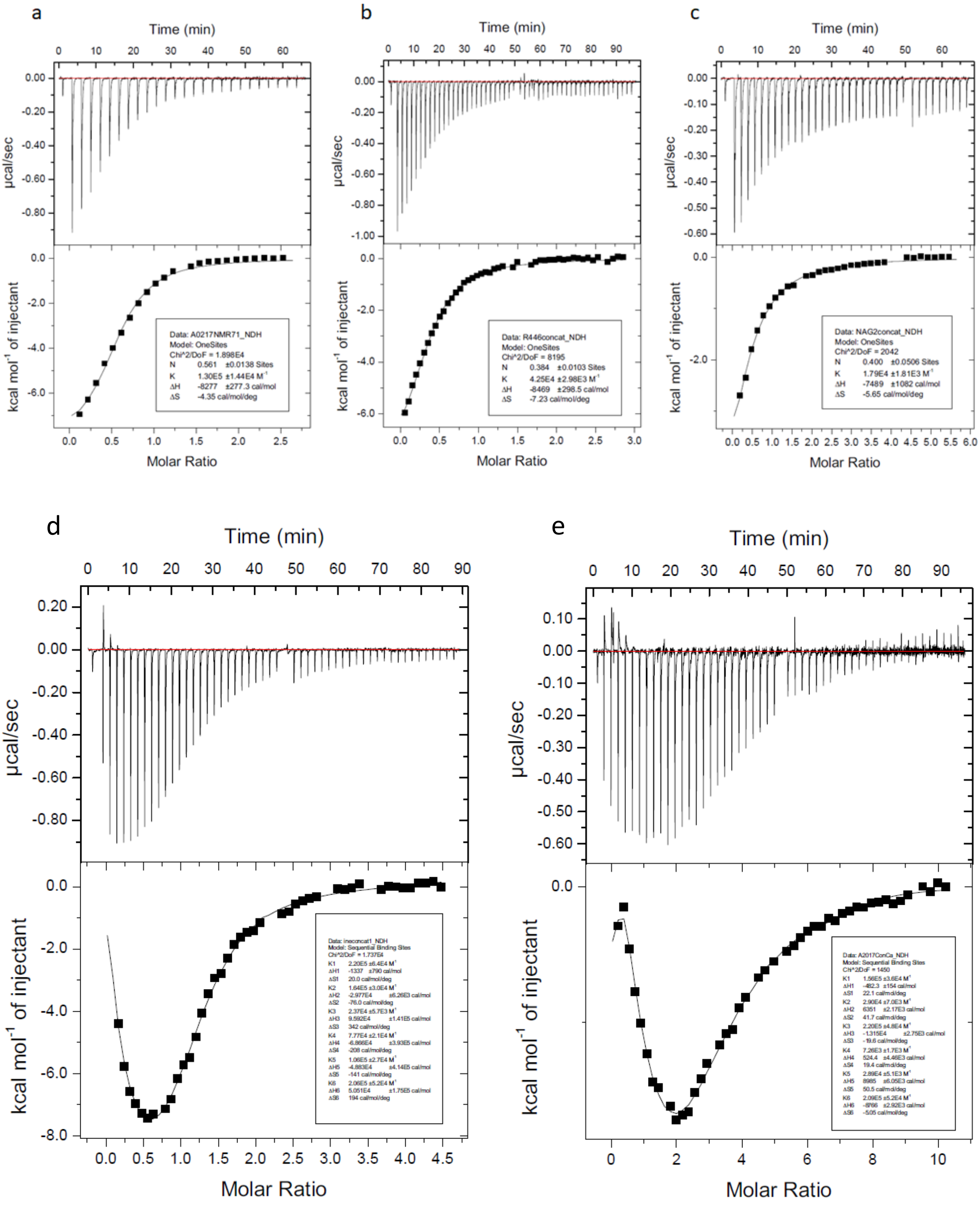


**Figure S7:** ArgF ITC titrations for NMR711 (a), and NMR446 (b), NAG (c), L-arginine (d), and L-canavanine (e).

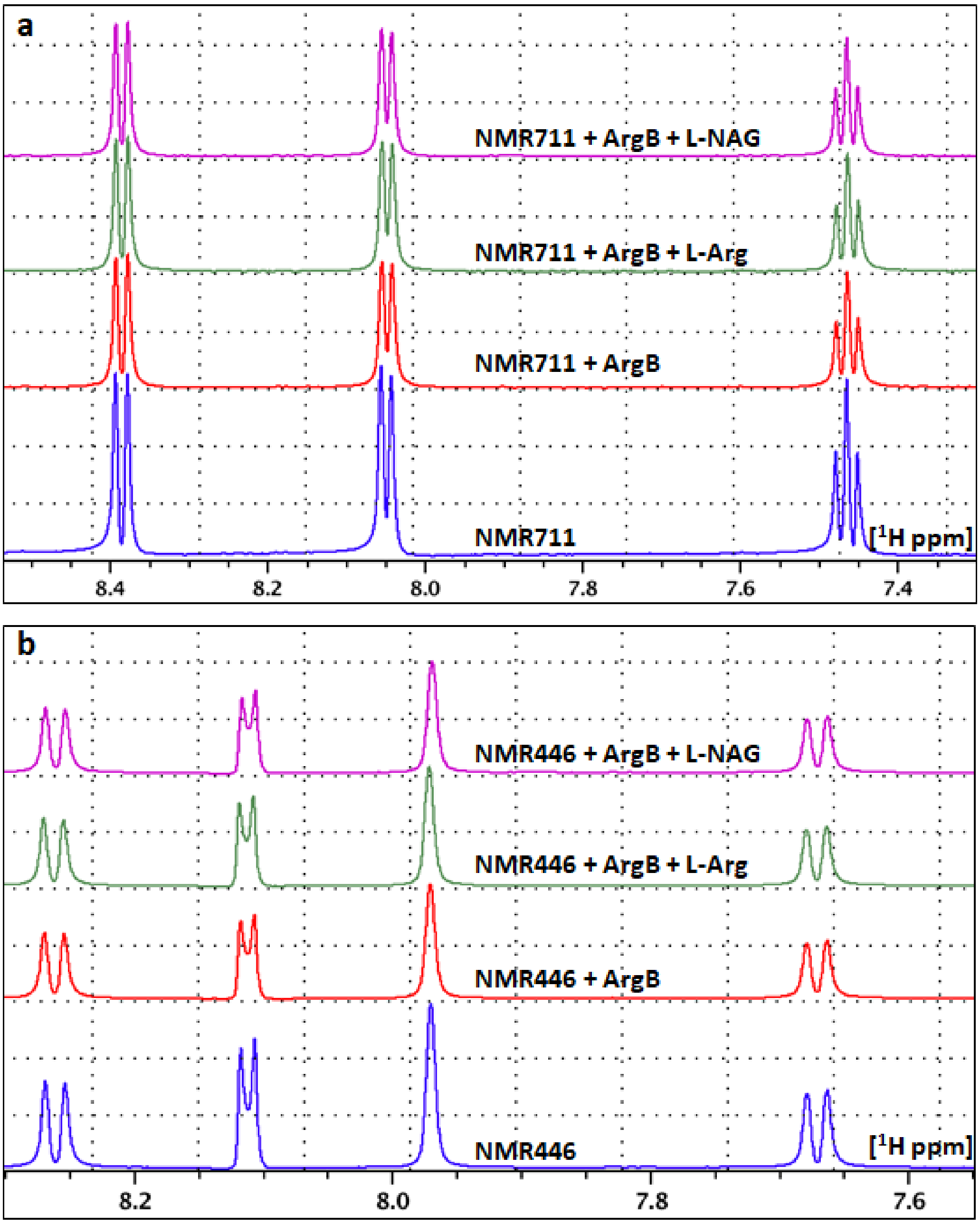


**Figure S8:** Ligand-observed NMR (CPMG) technique to illustrate binding of compounds (A) NMR711 and (B) NMR446 to ArgB. $^{1}$H NMR spectra of the fragment in the absence (blue) and presence of ArgB (red) are shown, where a drop in overall signal in the presence of ArgB indicates ligand binding to the protein. $^{1}$H NMR spectra of fragment in the presence L-arginine (green) or L-NAG (magenta) are also overlaid, where no significant change in signal indicates that the fragments bind to an allosteric site.

**Table S2:** ArgB % inhibition at 2.5 mM.

| Compound | Fragment structure | % inhibition |
|---|---|---|
| NMR026 | 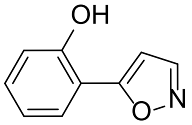 | 8 ± 1 |
| NMR043 | 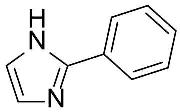 | 5 ± 2 |
| NMR078 | 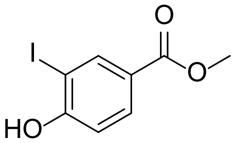 | 6 ± 1 |
| NMR082 | 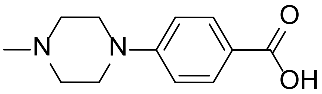 | 8 ± 1 |
| NMR314 | 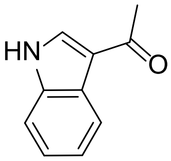 | 6 ± 3 |
| NMR323 | 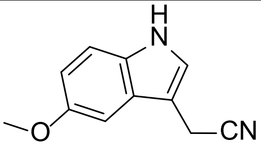 | 18 ± 2 |
| NMR462 | 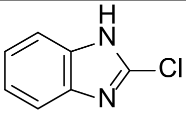 | 6 ± 1 |
| NMR469 | 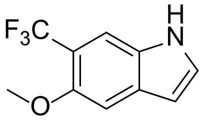 | 0 |
| NMR582 | 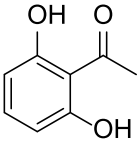 | 7 ± 3 |
| NMR612 | 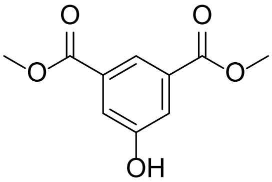 | 11 ± 3 |
| NMR617 | 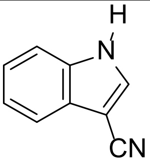 | 0 |

**Table S3:** X-ray crystallography data collection and final refinement statistics

| Protein | ArgB | ArgB | ArgB | ArgB | ArgB | ArgB | ArgB | ArgB | ArgB |
|---|---|---|---|---|---|---|---|---|---|
| Ligand# | **APO** | **NAG** | **L-arginine** | **L-canavanine** | **NMR711** | **NMR446** | **NMR026** | **NMR043** | **NMR078** |
| PDB ID | 7NLF | 7NLN | 7NLO | 7NLP | 7NNB | 7NLX | 7NLQ | 7NLR | 7NLS |
| **Data collection*** | | | | | | | | | |
| Space group | *R32* | $P6_3$ | *R32* | *R32* | *R32* | *R32* | *R32* | *R32* | *R32* |
| Cell parameters: | | | | | | | | | |
| a [Å] | 173.95 | 100.24 | 175.30 | 174.06 | 173.88 | 174.48 | 174.93 | 173.82 | 174.24 |
| b [Å] | 173.95 | 100.24 | 175.30 | 174.06 | 173.88 | 174.48 | 174.93 | 173.82 | 174.24 |
| c [Å] | 70.29 | 124.53 | 70.54 | 71.15 | 71.91 | 72.54 | 72.90 | 72.43 | 72.03 |
| α/β/γ [˚] | 90/90/120 | 90/90/120 | 90/90/120 | 90/90/120 | 90/90/120 | 90/90/120 | 90/90/120 | 90/90/120 | 90/90/120 |
| Resolution range [Å] | 86.98 – 2.08 (2.19 – 2.08) | 86.81 – 1.92 (2.02 – 1.92) | 63.97 – 1.82 (1.92 – 1.82) | 87.03– 2.21 (2.33 – 2.21) | 52.00 – 2.19 (2.30 – 2.19) | 87.24 – 2.23 (2.35 – 2.23) | 65.70 – 2.50 (2.63 – 2.50) | 86.91 – 2.25 (2.39 – 2.25) | 87.12 – 2.65 (2.79 – 2.65) |
| No. of observations | | | | | | | | | |
| total | 254931 (38396) | 566363 (40764) | 372764 (54564) | 154874 (20116) | 195399 (28703) | 156940 (20049) | 128210 (18834) | 199430 (31714) | 124133 (18498) |
| unique | 25346 (3662) | 54154 (7879) | 37098 (5393) | 20426 (2811) | 21590 (3120) | 20098 (2605) | 14890 (2140) | 19795 (3139) | 12283 (1774) |
| $R_{merge}$ | 0.045(0.876) | 0.071 (0.595) | 0.041 (0.975) | 0.057 (0.494) | 0.037 (0.787) | 0.041 (0.824) | 0.078 (0.863) | 0.047 (0.816) | 0.072 (1.213) |
| I/σ(I) | 22.2 (2.3) | 19.6 (2.5) | 23.9 (2.6) | 14.6 (2.2) | 27.7 (2.5) | 22.3 (2.2) | 14.4 (2.2) | 23.1 (2.3) | 20.3 (1.9) |
| CC(1/2) | 1.000 (0.885) | 0.999 (0.813) | 0.999 (0.895) | 0.999 (0.951) | 1.000 (0.905) | 0.999 (0.803) | 0.998 (0.747) | 0.999 (0.843) | 0.999 (0.759) |
| Completeness [%] | 100 (99.9) | 99.9 (99.6) | 100 (99.9) | 98.9 (94.0) | 100 (100) | 97.7 (87.3) | 99.8 (99.3) | 99.8 (98.1) | 100 (100) |
| Multiplicity | 10.1 (10.5) | 10.5 (5.2) | 10.0 (10.1) | 7.6 (7.2) | 9.1 (9.2) | 7.8 (7.7) | 8.6 (8.8) | 10.1 (10.1) | 10.1 (10.4) |
| **Refinement** | | | | | | | | | |
| Refinement program | PHENIX | PHENIX | PHENIX | PHENIX | PHENIX | PHENIX | PHENIX | PHENIX | PHENIX |
| Resolution [Å] | 86.98 – 2.08 | 86.81 – 1.92 | 63.97 – 1.82 | 87.03– 2.21 | 52.00 – 2.19 | 87.24 – 2.23 | 65.70 – 2.50 | 86.91 – 2.25 | 87.12 – 2.65 |
| No. reflections | 25346 | 54154 | 37098 | 20426 | 21590 | 20098 | 14890 | 19795 | 12283 |
| $R_{work}/R_{free}$ [%] | 18.2/21.6 | 16.5/19.6 | 18.8/20.3 | 20.5/24.1 | 19.9/25.9 | 19.5/23.7 | 18.7/22.5 | 18.4/21.4 | 19.3/25.6 |
| RMS deviations | | | | | | | | | |
| Bonds [Å] | 0.008 | 0.008 | 0.007 | 0.008 | 0.009 | 0.009 | 0.009 | 0.008 | 0.009 |
| Angles [˚] | 0.881 | 1.016 | 0.859 | 1.014 | 0.978 | 0.963 | 0.973 | 0.908 | 1.093 |
| Ramachandran | | | | | | | | | |
| Favoured [%] | 97 | 97 | 97 | 96 | 95 | 97 | 96 | 97 | 95 |
| Outliers [%] | 0.3 | 0.3 | 0.3 | 0.3 | 0.3 | 0.7 | 1.0 | 1.0 | 1.0 |

* Parameters shown in brackets are for the highest resolution shell

**Table S3:** X-ray crystallography data collection and final refinement statistics (cont)

| Protein | **ArgB** | **ArgB** | **ArgB** | **ArgB** | **ArgB** | **ArgB** | **ArgB** | **ArgB** | **ArgC** |
|---|---|---|---|---|---|---|---|---|---|
| Ligand# | **NMR082** | **NMR314** | **NMR323** | **NMR462** | **NMR469** | **NMR582** | **NMR612** | **NMR617** | **Apo** |
| PDB ID | 7NLT | 7NLU | 7NLW | 7NLY | 7NLZ | 7NM0 | 7NN7 | 7NN8 | 7NNI |
| **Data collection*** | | | | | | | | | |
| Space group | *R32* | *R32* | *R32* | *R32* | *R32* | *R32* | *R32* | *R32* | *C2* |
| Cell parameters: | | | | | | | | | |
| a [Å] | 173.63 | 174.49 | 174.35 | 174.94 | 174.06 | 174.35 | 174.16 | 174.65 | 140.69 |
| b [Å] | 173.63 | 174.49 | 174.35 | 174.94 | 174.06 | 174.35 | 174.16 | 174.65 | 77.99 |
| c [Å] | 71.99 | 71.73 | 72.79 | 72.96 | 72.81 | 72.1 | 72.06 | 72.12 | 88.29 |
| α/β/γ [˚] | 90/90/120 | 90/90/120 | 90/90/120 | 90/90/120 | 90/90/120 | 90/90/120 | 90/90/120 | 90/90/120 | 90/127.5/90 |
| Resolution range [Å] | 50.12 – 2.25 (2.35 – 2.23) | 64.80 – 2.24 (2.36 – 2.24) | 87.18 – 2.32 (2.45 – 2.32) | 87.47 – 2.25 (2.37 – 2.25) | 87.03 – 2.16 (2.28 – 2.16) | 87.17 – 2.28 (2.40 – 2.28) | 65.02 – 2.17 (2.29 – 2.17) | 87.32 – 2.27 (2.39 – 2.27) | 68.45 – 1.54 (1.63 – 1.54) |
| No. of observations | | | | | | | | | |
| total | 209659 (30760) | 180087 (26623) | 157958 (24489) | 155612 (23619) | 200575 (29524) | 170194 (25837) | 193857 (25606) | 176391 (26299) | 545085 (74721) |
| unique | 20440 (2950) | 20234 (2929) | 18318 (2657) | 20451 (2982) | 22616 (3285) | 18982 (2771) | 21728 (2958) | 19599 (2834) | 110883 (16103) |
| $R_{merge}$ | 0.061 (0.947) | 0.047 (0.946) | 0.066 (1.137) | 0.051 (0.845) | 0.053 (0.862) | 0.045 (0.893) | 0.053 (0.804) | 0.047 (0.960) | 0.191 (1.405) |
| I/σ(I) | 18.0 (2.2) | 20.2 (2.4) | 14.8 (1.9) | 17.4 (2.1) | 17.8 (2.1) | 22.3 (2.2) | 19.1 (2.3) | 20.6 (2.1) | 5.4 (1.3) |
| CC(1/2) | 0.999 (0.866) | 0.999 (0.900) | 0.999 (0.754) | 0.999 (0.706) | 0.999 (0.808) | 0.999 (0.892) | 0.999 (0.839) | 0.999 (0.796) | 0.986 (0.465) |
| Completeness [%] | 100 (100) | 99.8 (100) | 99.7 (100) | 100 (100) | 100 (100) | 99.0 (100) | 98.3 (93) | 100 (100) | 99.9 (99.9) |
| Multiplicity | 10.3 (10.4) | 8.9 (9.1) | 8.6 (9.2) | 7.6 (7.9) | 8.9 (9.0) | 9.0 (9.3) | 8.9 (8.7) | 9.0 (9.3) | 4.9 (4.6) |
| **Refinement** | | | | | | | | | |
| Refinement program | PHENIX | PHENIX | PHENIX | PHENIX | PHENIX | PHENIX | PHENIX | PHENIX | PHENIX |
| Resolution [Å] | 50.12 – 2.25 | 64.80 – 2.24 | 87.18 – 2.32 | 87.47 – 2.25 | 87.03 – 2.16 | 87.17 – 2.28 | 65.02 – 2.17 | 87.32 – 2.27 | 68.45 – 1.54 |
| No. reflections | 20440 | 20234 | 18318 | 20451 | 22616 | 18982 | 21728 | 19599 | 110292 |
| $R_{work}/R_{free}$ [%] | 19.0/25.1 | 19.5/23.6 | 21.1/24.3 | 19.2/23.9 | 19.1/23.8 | 19.3/22.5 | 18.8/23.6 | 18.9/23.3 | 19.3/21.2 |
| RMS deviations | | | | | | | | | |
| Bonds [Å] | 0.008 | 0.009 | 0.009 | 0.009 | 0.009 | 0.008 | 0.009 | 0.009 | 0.010 |
| Angles [˚] | 0.982 | 1.111 | 1.100 | 1.021 | 0.975 | 0.982 | 0.965 | 1.037 | 1.36 |
| Ramachandran | | | | | | | | | |
| Favoured [%] | 97 | 96 | 94 | 96 | 97 | 96 | 97 | 95 | 98.7 |
| Outliers [%] | 0.3 | 0.3 | 0.7 | 0.3 | 0.3 | 0.3 | 0.3 | 0.7 | 0 |

* Parameters shown in brackets are for the highest resolution shell

**Table S3:** X-ray crystallography data collection and final refinement statistics (cont)

| Protein | **ArgC** | **ArgC** | **ArgC** | **ArgC** | **ArgC** | **ArgD** | **ArgD** | **ArgD** | **ArgF** |
|---|---|---|---|---|---|---|---|---|---|
| Ligand# | **NADP** | **NMR322** | **NMR401** | **NMR571** | **NMR863** | **PLP** | **NMR608** | **NMR868** | **Apo** |
| PDB ID | 7NNQ | 7NOT | 7NPJ | 7NNR | 7NPH | 7NN1 | 7NN4 | 7NNC | 7NNF |
| **Data collection*** | | | | | | | | | |
| Space group | *P2₁* | *P2₁* | *P2₁* | *C2* | *P2₁* | *P2₁* | *P2₁* | *P2₁* | *P2₁* |
| Cell parameters: | | | | | | | | | |
| a [Å] | 69.05 | 83.40 | 84.40 | 140.71 | 84.28 | 65.06 | 65.19 | 65.46 | 91.34 |
| b [Å] | 133.83 | 132.00 | 132.85 | 78.23 | 132.79 | 184.26 | 184.35 | 184.94 | 142.38 |
| c [Å] | 83.05 | 124.55 | 122.71 | 87.64 | 122.83 | 71.54 | 71.30 | 71.58 | 98.28 |
| α/β/γ [˚] | 90/109.4/90 | 90/97.06/90 | 90/90.06/90 | 90/127.51/90 | 90/90.17/90 | 90/106.1/90 | 90/106.21/90 | 90/106.1/90 | 90/117.2/90 |
| Resolution range [Å] | 55.52 – 1.73 (1.77 – 1.73) | 63.91 – 2.54 (2.61 – 2.54) | 71.24 – 2.81 (2.88 – 2.81) | 69.51 – 1.7 (1.73 – 1.7) | 69.59 – 2.57 (2.64 – 2.57) | 62.50 – 1.54 (1.62 – 1.54) | 68.47 – 1.47 (1.51 – 1.47) | 68.78 – 1.70 (1.73 – 1.70) | 81.12 – 1.52 (1.61 – 1.52) |
| No. of observations | | | | | | | | | |
| total | 804727 (58410) | 463192 (36392) | 353903 (26408) | 417201 (15597) | 466260 (34746) | 1065135 (75420) | 1192624 (51854) | 948793 (41174) | 1731990 (209268) |
| unique | 145458 (10386) | 87548 (6486) | 65900 (4807) | 81979 (3945) | 86043 (6338) | 218441 (20903) | 265823 (17510) | 177420 (8457) | 338738 (49398) |
| $R_{merge}$ | 0.048 (1.275) | 0.206 (2.135) | 0.117 (1.641) | 0.031 (0.091) | 0.067 (1.473) | 0.068 (0.873) | 0.035 (0.693) | 0.027 (0.082) | 0.096 (0.978) |
| I/σ(I) | 13.9 (1.0) | 7.3 (0.8) | 6.9 (1.0) | 29.3 (10.2) | 11.0 (1.0) | 9.4 (1.1) | 16.5 (1.2) | 34.0 (12.1) | 7.4 (1.0) |
| CC(1/2) | 0.999 (0.462) | 0.986 (0.317) | 0.993 (0.321) | 0.999 (0.988) | 0.999 (0.522) | 0.998 (0.438) | 0.999 (0.500) | 0.999 (0.994) | 0.994 (0.449) |
| Completeness [%] | 98.4 (95.4) | 99.3 (99.9) | 99.9 (100) | 99 (89.6) | 100 (100) | 91.7 (60.2) | 97.4 (86.7) | 99.3 (95.6) | 99.9 (100.0) |
| Multiplicity | 5.5 (5.6) | 5.3 (5.6) | 5.4 (5.5) | 5.1 (4) | 5.4 (5.5) | 4.9 (3.6) | 4.5 (3.0) | 5.3 (4.9) | 5.1 (4.2) |
| **Refinement** | | | | | | | | | |
| Refinement program | PHENIX | PHENIX | PHENIX | PHENIX | PHENIX | PHENIX | PHENIX | PHENIX | PHENIX |
| Resolution [Å] | 45.09 – 1.73 | 58.44 – 2.54 | 69.46 – 2.81 | 40.57 – 1.7 | 69.4 – 2.57 | 46.81 – 1.539 | 64.18 – 1.47 | 45.91 – 1.70 | 81.12 – 1.52 |
| No. reflections | 145435 | 87492 | 65781 | 81962 | 85901 | 218262 | 265761 | 177364 | 338688 |
| $R_{work}/R_{free}$ [%] | 19.1/21.1 | 21.5/27.5 | 25.7/32.7 | 15.8/17.5 | 26/31.1 | 18/20 | 18/19.6 | 14.7/16.3 | 16.9/18.3 |
| RMS deviations | | | | | | | | | |
| Bonds [Å] | 0.008 | 0.011 | 0.009 | 0.006 | 0.010 | 0.009 | 0.008 | 0.009 | 0.007 |
| Angles [˚] | 1.27 | 1.5 | 1.27 | 0.84 | 1.46 | 1.44 | 1.41 | 1.2 | 1.093 |
| Ramachandran | | | | | | | | | |
| Favoured [%] | 97.9 | 96 | 86.99 | 98.8 | 93.1 | 97.6 | 97.60 | 97.5 | 98 |
| Outliers [%] | 0.07 | 0.3 | 2.08 | 0 | 0.7 | 0.2 | 0.3 | 0.4 | 0.4 |

* Parameters shown in brackets are for the highest resolution shell

**Table S3:** X-ray crystallography data collection and final refinement statistics (cont)

| Protein | ArgF | ArgF | ArgF | ArgF | ArgF | ArgF | ArgF | ArgF | ArgF |
|---|---|---|---|---|---|---|---|---|---|
| Ligand# | **CP** | **NMR007** | **NMR078** | **NMR288** | **NMR464** | **NMR502** | **NMR801** | **NMR812** | **NMR817** |
| PDB ID | 7NNV | 7NNZ | 7NNW | 7NNY | 7NOS | 7NOR | 7NP0 | 7NOU | 7NOV |
| **Data collection*** | | | | | | | | | |
| Space group | $P2_1$ | $P2_1$ | $P2_1$ | $P2_1$ | $P2_1$ | $P2_1$ | $P2_1$ | $P2_1$ | $P2_1$ |
| Cell parameters: | | | | | | | | | |
| a [Å] | 91.44 | 91.62 | 91.80 | 91.95 | 92.00 | 91.45 | 92.04 | 92.30 | 92.26 |
| b [Å] | 142.37 | 143.00 | 143.51 | 143.79 | 143.61 | 142.73 | 143.45 | 144.66 | 144.42 |
| c [Å] | 99.02 | 97.59 | 97.52 | 97.28 | 97.82 | 97.70 | 97.53 | 97.76 | 97.35 |
| α/β/γ [˚] | 90/117.2/90 | 90/117.6/90 | 90/117.7/90 | 90/117.6/90 | 90/117.5/90 | 90/117.5/90 | 90/117.5/90 | 90/117.8/90 | 90/117.8/90 |
| Resolution range [Å] | 88.10 – 1.67 (1.76 – 1.67) | 81.20 – 1.68 (1.77 – 1.68) | 86.36 – 1.78 (1.87 – 1.78) | 86.21 – 1.57 (1.66 – 1.57) | 81.61 – 1.77 (1.86 – 1.77) | 81.12 – 1.59 (1.68 – 1.59) | 70.46 – 1.76 (1.85 – 1.76) | 86.47 – 1.98 (2.08 – 1.98) | 86.15 – 1.90 (2.00 – 1.90) |
| No. of observations | | | | | | | | | |
| total | 1330397 (151816) | 1305874 (162377) | 1131278 (166972) | 1448621 (105414) | 1170700 (168321) | 1416962 (114917) | 1149709 (167284) | 1604954 (230134) | 840668 (80109) |
| unique | 259225 (36285) | 252690 (36825) | 214281 (31302) | 294032 (31609) | 219906 (32044) | 271697 (26818) | 222029 (32413) | 157930 (23055) | 170994 (20551) |
| $R_{merge}$ | 0.051 (1.051) | 0.089 (0.898) | 0.078 (1.140) | 0.043 (0.937) | 0.045 (1.140) | 0.052 (1.109) | 0.047 (0.508) | 0.083 (0.895) | 0.057 (0.312) |
| I/σ(I) | 13.6 (0.9) | 9.3 (1.7) | 10.9 (1.1) | 15.4 (0.9) | 15.5 (1.1) | 13.0 (0.9) | 16.0 (2.7) | 16.1 (2.7) | 14.1 (2.6) |
| CC(1/2) | 0.999 (0.460) | 0.997 (0.456) | 0.999 (0.530) | 0.999 (0.424) | 0.999 (0.551) | 0.999 (0.427) | 0.999 (0.801) | 0.998 (0.830) | 0.998 (0.884) |
| Completeness [%] | 99.2 (95.2) | 99.9 (99.8) | 99.9 (100) | 94.6 (69.8) | 100 (100) | 92.2 (62.3) | 99.9 (100) | 99.9 (100) | 96.7 (79.6) |
| Multiplicity | 5.1 (4.2) | 5.2 (4.4) | 5.3 (5.3) | 4.9 (3.3) | 5.3 (5.3) | 5.2 (4.3) | 5.2 (5.2) | 10.2 (10.0) | 4.9 (3.9) |
| **Refinement** | | | | | | | | | |
| Refinement program | PHENIX | PHENIX | PHENIX | PHENIX | PHENIX | PHENIX | PHENIX | PHENIX | PHENIX |
| Resolution [Å] | 88.10 – 1.67 | 81.20 – 1.68 | 86.36 – 1.78 | 86.21 – 1.57 | 81.61 – 1.77 | 81.12 – 1.59 | 70.46 – 1.76 | 86.47 – 1.98 | 86.15 – 1.90 |
| No. reflections | 259158 | 252638 | 423145 | 293962 | 219865 | 271656 | 221982 | 157873 | 170957 |
| $R_{work}/R_{free}$ [%] | 16.3/18.0 | 16.7/18.3 | 18.1/20.6 | 16.6/18.4 | 17.0/19.2 | 16.7/18.6 | 16.2/18.2 | 17.4/20.1 | 16.1/19.0 |
| RMS deviations | | | | | | | | | |
| Bonds [Å] | 0.009 | 0.010 | 0.009 | 0.007 | 0.012 | 0.008 | 0.007 | 0.011 | 0.009 |
| Angles [˚] | 1.463 | 1.622 | 1.419 | 1.213 | 1.596 | 1.177 | 0.959 | 1.572 | 1.483 |
| Ramachandran | | | | | | | | | |
| Favoured [%] | 97 | 97 | 97 | 98 | 97 | 98 | 97 | 97 | 97 |
| Outliers [%] | 0.4 | 0.4 | 0.3 | 0.3 | 0.4 | 0.4 | 0.3 | 0.7 | 0.4 |

* Parameters shown in brackets are for the highest resolution shell
